# KDM7B-mediated demethylation of RNF113A regulates small cell lung cancer sensitivity to alkylation damage

**DOI:** 10.64898/2026.01.03.697470

**Authors:** Tanveer Ahmad, Xiaojie Yang, Anne-Emmanuelle Foucher, Lingnan Ren, Ning Tsao, Natasha Flores, Jinkai Wan, Lucid Belmudes, Etienne Dubiez, Monika Chandan Bhowmik, Jessica Vayr, Simone Hausmann, Florent Chuffart, Xiaoyin Lu, Sandrine Blanchet, Tourkian Chasan, Faycal Boussouar, Yohann Couté, Nima Mosammaparast, Fei Lan, Jan Kadlec, Pawel K. Mazur, Nicolas Reynoird

**Affiliations:** Grenoble Alpes University, INSERM U1209, CNRS UMR 5309, Institut pour l’Avancée des Biosciences (IAB), ProMeDy, 38000 GRENOBLE, France; Department of Experimental Radiation Oncology, The University of Texas MD Anderson Cancer Center, Houston, Texas; Grenoble Alpes University, CNRS, CEA, IBS, 38000 Grenoble, France; Shanghai Key Laboratory of Medical Epigenetics, International Co-laboratory of Medical Epigenetics and Metabolism, Ministry of Science and Technology, Institutes of Biomedical Sciences, Key Laboratory of Carcinogenesis and Cancer Invasion, Ministry of Education, Liver Cancer Institute, Zhongshan Hospital, Fudan University, Shanghai, China; Department of Pathology and Immunology and Department of Medicine, Center for Genome Integrity, Washington University in St. Louis School of Medicine, St. Louis, Missouri; Grenoble Alpes University, CEA, INSERM, UA13 BGE, CNRS CEA FR2048, 38000 Grenoble, France

## Abstract

Chemoresistance remains a major obstacle to effective cancer treatment, often driven by enhanced DNA repair mechanisms that enable tumor cells to withstand genotoxic therapies. One such pathway involves the atypical DNA damage repair complex ALKBH3–ASCC, activated by the E3 ligase RNF113A in response to alkylation damage. We previously showed that SMYD3-dependent methylation of RNF113A stimulates this pathway, enhancing DNA repair and promoting resistance.

Here, we identify KDM7B/PHF8 as the bona fide RNF113A demethylase, establishing one of the first functional examples of a dynamic, reversible non-histone methylation event regulating genome integrity. KDM7B antagonizes SMYD3 activity by maintaining low levels of methylated RNF113A, thereby limiting ASCC activation and sensitizing cancer cells to alkylating agents. To dissect this regulation in depth, we focused on small cell lung cancer (SCLC), a particularly aggressive malignancy characterized by limited therapeutic options and rapid acquisition of resistance. In SCLC, high KDM7B levels correlate with improved patient prognosis, whereas xenografts with reduced expression exhibit diminished responses to alkylating treatment. Moreover, CRISPR-based on/off modulation of KDM7B in genetically engineered SCLC mouse models demonstrates its central role in determining tumor response to chemotherapy.

Our findings position the RNF113A–ASCC axis as a central modulator of chemoresistance, regulated through a post-translational methylation switch representing an innovative therapeutic vulnerability that could be exploited to enhance the efficacy of alkylating agents. Targeting this pathway may provide new opportunities to overcome chemoresistance, with KDM7B levels serving as a predictive biomarker to guide treatment in SCLC.

## INTRODUCTION

Chemoresistance remains a central obstacle to effective cancer treatment, limiting the long-term benefits of both cytotoxic and targeted therapies [1]. A key driver of resistance is the ability of tumor cells to engage DNA damage repair (DDR) pathways that mitigate the cytotoxicity of genotoxic agents [2,3]. Among commonly used therapies, platinum-based and alkylating chemotherapies remain widely used across multiple cancer types, yet their efficacy is consistently undermined by the emergence of resistance. Dissecting the molecular circuits that underlie DNA repair–mediated chemoresistance therefore represents a critical step toward improving clinical outcomes, especially in cancer with limited therapeutic options.

Alkylation damage can be repaired through multiple mechanisms, including base excision repair initiated by alkyladenine DNA glycosylase (AAG), direct reversal via O^6-methylguanine DNA methyltransferase (MGMT), and oxidative demethylation by AlkB family dioxygenases [4]. More recently, an atypical signaling pathway has been described in which the E3 ubiquitin ligase RNF113A links the ASCC (activating signal cointegrator complex) to ALKBH3, coordinating RNA and DNA alkylation repair [5]. We previously showed that RNF113A is methylated by the lysine methyltransferase SMYD3, a post-translational modification (PTM) that enhances its ligase activity, boosts ASCC–ALKBH3 function, and promotes cellular tolerance to alkylating agents [6]. This RNF113A methylation switch thus represents a critical mechanism coupling DDR efficiency to chemoresistance.

To explore the importance of this mechanism in a clinically relevant context, we focused on small cell lung cancer (SCLC), one of the most lethal solid malignancies. Lung cancer comprises two major histological subtypes: non-small cell lung cancer (NSCLC), which represents approximately 85% of lung cancer cases, and small cell lung cancer (SCLC), which accounts for the remaining 15% [7]. While therapeutic advances, including targeted therapies and immune checkpoint inhibitors, have significantly improved survival outcomes in NSCLC, progress in the treatment of SCLC has remained limited. Characterized by early metastasis and an exceptionally high proliferation rate and rapid acquisition of resistance, SCLC exhibits one of the highest mortality rates across all cancer types, making it a prototypical model of therapy-refractory cancer [8].

This therapeutic stagnation can be partly attributed to the absence of clearly defined, druggable genetic drivers and to date SCLC continues to be treated primarily with conventional chemotherapy. SCLC is initially highly responsive to cytotoxic agents, exceeding 60% overall response rates to first-line treatment in extensive-stage disease. However, these responses are almost universally transient. Most patients relapse within months, and recurrent disease is typically resistant to further treatment, resulting in a median survival of approximately 1 year in extensive-stage SCLC [8]. First-line platinum-based chemotherapy (cisplatin or carboplatin with etoposide) has replaced earlier alkylation-based regimens (e.g., cyclophosphamide + doxorubicin + vincristine) due to reduced toxicity, but offers no significant improvement in efficacy [9–11]. Alkylating agents retain some activity in platinum-refractory disease, though the reverse is rarely true, underscoring the unique therapeutic relevance of alkylation tolerance [11]. Recent advances in elucidating SCLC subtypes and the molecular circuitry of SCLC provide a promising foundation [12], but actionable insights remain sparse. Among other potential strategies, dissecting the cellular mechanisms that drive alkylation tolerance and chemoresistance represents a critical avenue toward improving outcomes for SCLC patients.

Given the lack of clearly defined genetic drivers in SCLC, increasing attention has turned toward regulatory processes that enable rapid adaptation to therapy. Post-translational modifications (PTMs) are among the most versatile of these processes, acting as reversible molecular switches that fine-tune protein activity in stress responses. Among the many PTMs, lysine methylation has drawn increasing interest: initially studied in the context of histone regulation, recent evidence suggests that this modification also extends to non-histone proteins, with the potential to influence DNA repair and therapy resistance [13]. As described earlier, we notably found that the methylation of RNF113A by the lysine methyltransferase SMYD3 promotes chemoresistance[6]. While lysine methylation was once considered irreversible, it is now well established that these marks are dynamically regulated on histones thanks to two major classes of demethylases: amine oxidases (e.g., KDM1A/B) and Jumonji C domain– containing proteins (JMJDs), comprising over 30 members [14,15]. However, the specific reversibility of lysine methylation on non-histone proteins remains poorly characterized, even though enzymes such as KDM1A act on a broad range of substrates [16,17]. Thus, the extent and functional relevance of dynamic methylation on non-histone targets remains an emerging and compelling area of investigation.

In this context, we investigated whether RNF113A methylation is reversible, and whether this influences dealkylation repair and chemoresistance in SCLC. We identified KDM7B, also known as Plant Homologous domain Finger protein 8 (PHF8), as a bona fide RNF113A demethylase. KDM7B is a Jumonji domain– containing demethylase that recognizes H3K4me3 via its PHD domain and removes repressive histone marks including H3K9me1/2, H3K27me1, and H4K20me1 [18]. Through these activities, KDM7B has been implicated in chromatin remodeling, transcriptional activation, and regulation of cell fate decisions in development and cancer [19–22]. Importantly, KDM7B has also been associated with DNA damage responses. Notably, its homolog in C. elegans modulates homologous recombination (HR) repair of double-strand breaks [23]. In addition, KDM7B preserves genome stability by regulating TOPBP1, which facilitates ATR activation under replication stress [24]. Finally, KDM7B promotes transcriptional recovery after DNA double-strand break repair by removing repressive H3K9me2 marks [24].

Our findings uncover a novel role for KDM7B in alkylation damage repair and previously unrecognized layer of post-translational control over DNA repair, with implications both for overcoming chemoresistance in SCLC and for broader principles of methylation signaling in cancer biology.

## RESULTS

### KDM7B demethylates RNF113A

The E3 ligase RNF113A is actively engaged in alkylated DNA damage response [5]. We previously discovered that RNF113A is trimethylated on its lysine 20 by the methyltransferase SMYD3, and that this modification participates in resistance to alkylation-based chemotherapy in SCLC cancer cells [6]. In addition, we noticed that RNF113A is regulated by a dynamic phosphorylation switch, and that its inactivation by the phosphatase PP4 is blocked by RNF113A methylation [6]. We wondered whether, in addition to RNF113A phosphorylation, the K20 trimethylation mark might also be dynamic, with a potential demethylase at play. While histone demethylation is a well-documented process of chromatin regulation [15], there is still little evidence of active non-histone proteins demethylation besides the broad spectrum of KDM1A/LSD1 demethylase [17]. The possibility of a specific RNF113A demethylase could have intriguing consequences on SCLC patients’ sensitivity to alkylating chemotherapy.

To tackle this possibility, we monitored RNF113A methylation level in HeLa cells after incubation with 5-Carboxy-8-hydroxyquinoline (IOX1), a potent inhibitor of 2OG oxygenases including JmjC demethylases. We used engineered HeLa cells stably expressing HA-tagged RNF113A to enrich the protein and monitor its methylation level thanks to the previously characterized RNF113AK20me^3^ specific antibody. We observed a striking increase of methylated RNF113A levels upon IOX1 treatment while SMYD3 levels remained unaffected, suggesting that a JmjC demethylase is capable to counteract SMYD3 activity on RNF113A (Figure 1A).

**Figure 1:**
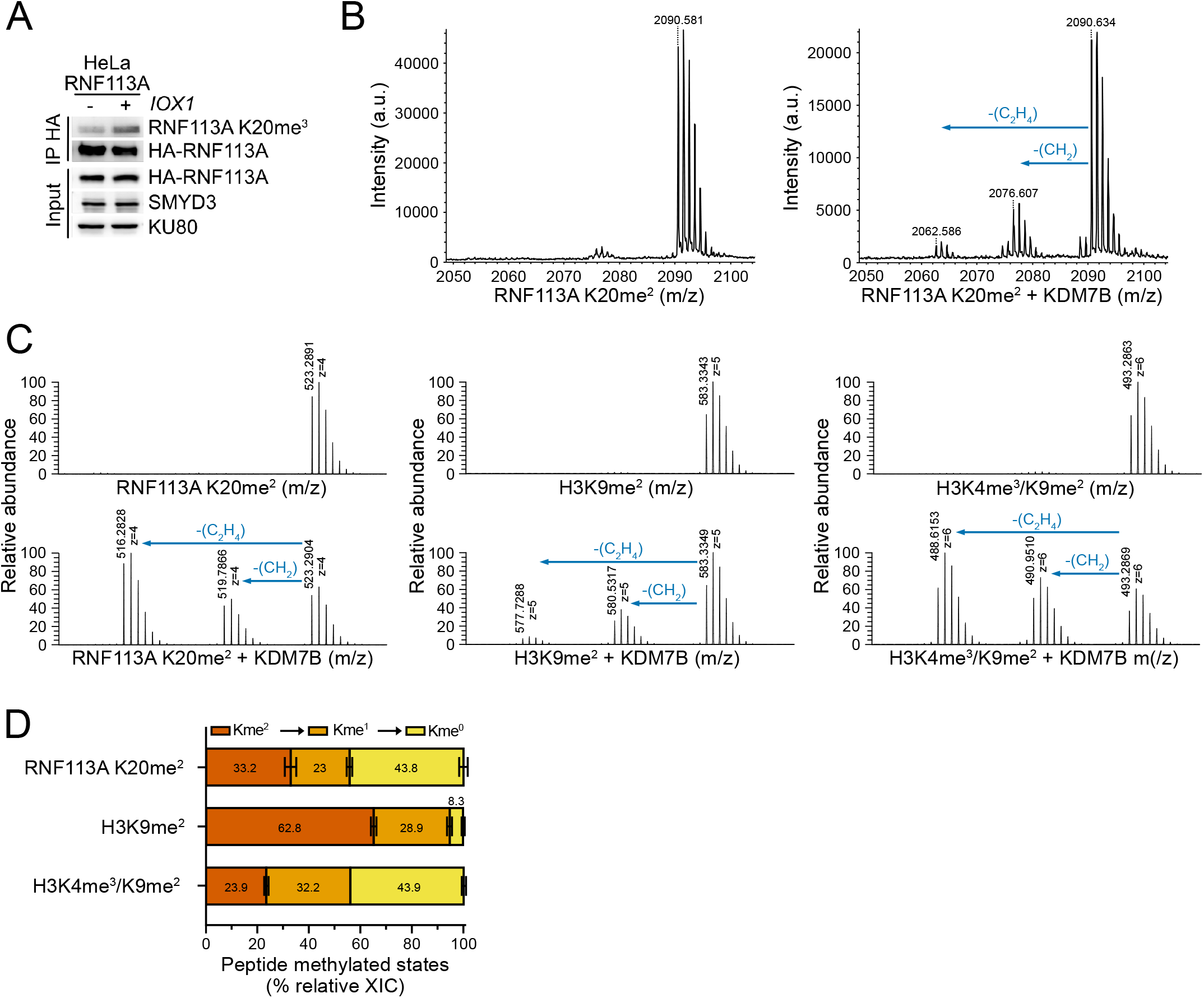
RNF113A is a genuine substrate of KDM7B. **A**, Immunodetection with indicated antibodies of methylated RNF113A levels after total RNF113A immunoprecipitation from HeLa cells stably expressing HA-RNF113A, either pre-treated with DMSO (Control) or treated with IOX1 inhibitor for 72 hours. KU80 was used as a loading control. **B**, Representative spectra after MALDI-mass spectrometry analyses of RNF113A *in vitro* demethylation assays using RNF113A K20me^2^ peptide in the absence (left panel) or presence (right panel) of the demethylase KDM7B. **C**, Representative spectra after ESI-mass spectrometry analyses of KDM7B-mediated demethylation of either RNF113A K20me^2^ (left panel), H3K9me^2^ (middle panel) and H3K4me^3^K9me^2^ (right panel) peptides. **D**, Quantification of the different states of methylation observed after KDM7B-mediated demethylation of RNF113A K20me^2^, H3K9me^2^ and H3K4me^3^K9me^2^. Each condition represents the mean with SEM of three technical replicates.

To identify the specific RNF113A demethylase, we performed a mass-spectrometry-based screen on a panel of recombinant demethylases representative of the different KDMs subfamilies [25] using either RNF113A lysine 20 mono, di or trimethylated peptides. Indeed, similar to lysine methyltransferases capable to catalyze a specific level of lysine methylation (mono, di or tri-methylation), demethylases with characterized activity on methylated histones show a preference for a given state of methylated lysines. We tested over 15 demethylases, including the amine oxidases KDM1A and KDM1B -even though they are not inhibited by IOX-1 - due to existing evidence of their broad range of non-histone protein targets [17]. Remarkably, we found a clear activity for a single demethylase, KDM7B, also known as PHF8, on methylated RNF113A (Figure 1B and Supplementary Figure 1A). KDM7B is a histone demethylase known as a transcriptional activator by erasing repressive histone marks, with preferential specificity for the H3K9me^2^ mark [18,21]. Of note, the PHD domain of KDM7B binds to H3K4me^3^ and this modification improves H3K9 demethylation efficiency. Interestingly, KDM7B is known to play context-dependent roles in cancer and among various functions, participates to DNA repair [24,26–28]. We noticed that KDM7B was more efficient on the dimethylated state of RNF113A (RNF113A K20me^2^, Supplementary Figure 1B), coherent with its preference for the lysine K9 dimethylated state of Histone H3 [18]. We then tested different recombinant KDM7B constructs and found that KDM7B 38-494 (numbering based on KDM7B isoform 1 Q9UPP1-1, and referred to as KDM7B short) purified from insect cells was highly active and preferentially used for further assays (Supplementary Figure 1C-D). We next compared KDM7B activity on methylated RNF113A and histone H3 and found that the enzyme demethylates RNF113A K20me^2^ in a similar efficacy than H3K4me^3^/K9me^2^ and significantly better than H3K9me^2^ (Figure 1C-D and Supplementary Figure 1E). In addition, we validated RNF113A K20me^2^ demethylation by KDM7B using a second indirect approach measuring succinate levels, a by-product of demethylation reaction (Succinate-Glo assay, Supplementary Figure 1F). Therefore, our results show that KDM7B efficiently demethylates RNF113A *in vitro*.

### KDM7B - RNF113A interaction study

Our observation that RNF113A is as efficiently demethylated by KDM7B as H3K4me^3^/H3K9me^2^ is particularly striking, given that RNF113A does not appear to possess an adjacent modification equivalent of the H3K4me^3^/H3K9me^2^ peptide recognized by the KDM7B PHD domain to strengthen the enzyme interaction with RNF113A. Thus, we aimed to better characterize the catalytic activity and binding of KDM7B on RNF113A. Using *in vitro* demethylation assays and mass spectrometry analysis, we calculated the KDM7B enzymatic velocity (units of demethylated substrates per unit of KDM7B catalyzed in one hour) for a given concentration of RNF113A and histone H3 peptides. We observed that KDM7B has a higher Michaelis-Menten constant (Km) for methylated RNF113A compared to methylated H3 (1.088 vs 0.326 μM) and a higher maximum velocity reaction (Vmax) (10.56 vs 2.63 u/h) (Figure 2A). This suggests that while KDM7B’s affinity for RNF113A K20me^2^ is lower compared to H3K4me^3^/K9me^2^ (higher Km), RNF113A K20me^2^ is a highly potent substrate of KDM7B (higher Vmax). We measured binding affinities between the catalytic domain of KDM7B (KDM7B short, see Supplementary Figure 1C) and different methylation states of RNF113A peptides by isothermal titration calorimetry (ITC). Consistent with our previous observations, we did not detect any interaction between KDM7B and unmethylated RNF113A peptide (Figure 2B). Interaction with a trimethylated lysine by KDM7B was previously shown to be incompatible with the binding of the α-ketoglutarate cofactor [18,29]. In agreement with this observation, although our results show that the trimethylated RNF113A peptide may interacts with KDM7B in the absence of α-ketoglutarate, the noisier binding profile prevented determination of a reliable *K*_*d*_ value (Supplementary Figure 2A). Conversely, RNF113A peptides bearing mono- and dimethylation at K20 (RNF113A-K20me^1^ and RNF113A-K20me^2^) unambiguously interacted with KDM7B with dissociation constants (*K*_*d*_) of 19.5 and 34 μM, respectively (Figure 2B and Supplementary Figure 2A). Of note, the measured affinities of KDM7B for RNF113A-K20me^1^ and RNF113A-K20me^2^ are lower than that for H3K4me^3^K9me^2^ (1 μM), but higher than that for H3K9me^2^ alone, previously reported to be negligible [18]. We confirmed these results through direct peptide pulldown assays of KDM7B, where we observed that dimethylated RNF113A interacts with KDM7B slightly better than H3K9me^2^ but less efficiently than H3K4me^3^/K9me^2^ (Figure 2C). The higher affinity of KDM7B for H3K4me^3^/K9me^2^ is attributed to the additional recognition of the H3K4me^3^ mark by the KDM7B PHD domain [18,29] (Figure 2D). To gain insight into the molecular details of RNF113A recognition by KDM7B, we used Alphafold3 structure prediction [30]. Alphafold3 predicts, with high confidence, contacts between the catalytic KDM7B Jmj domain and RNF113A only in the presence of methylation marks on RNF113A K20. In a model including the KDM7B JmjC domain (residues 114-482, see Supplementary Figure 1C) and RNF113A-K20me^2^ (residues 14-27), modified K20 inserts into the JmjC catalytic site (Figure 2E, Supplementary Figure 2B-C). The substrate peptide is predicted to form several side and main chain contacts between residues 18-24 and KDM7B, including salt bridge interactions of K21 with KDM7B D194 and D203 (Figure 2F). No contacts with the PHD domain were predicted with high confidence, consistent with the lower KDM7B affinity for RNF113A-K20me^2^ compared to H3K4me^3^/K9me^2^. Together, these results demonstrate that RNF113A-K20me^2^ is a genuine substrate of the KDM7B demethylase.

**Figure 2:**
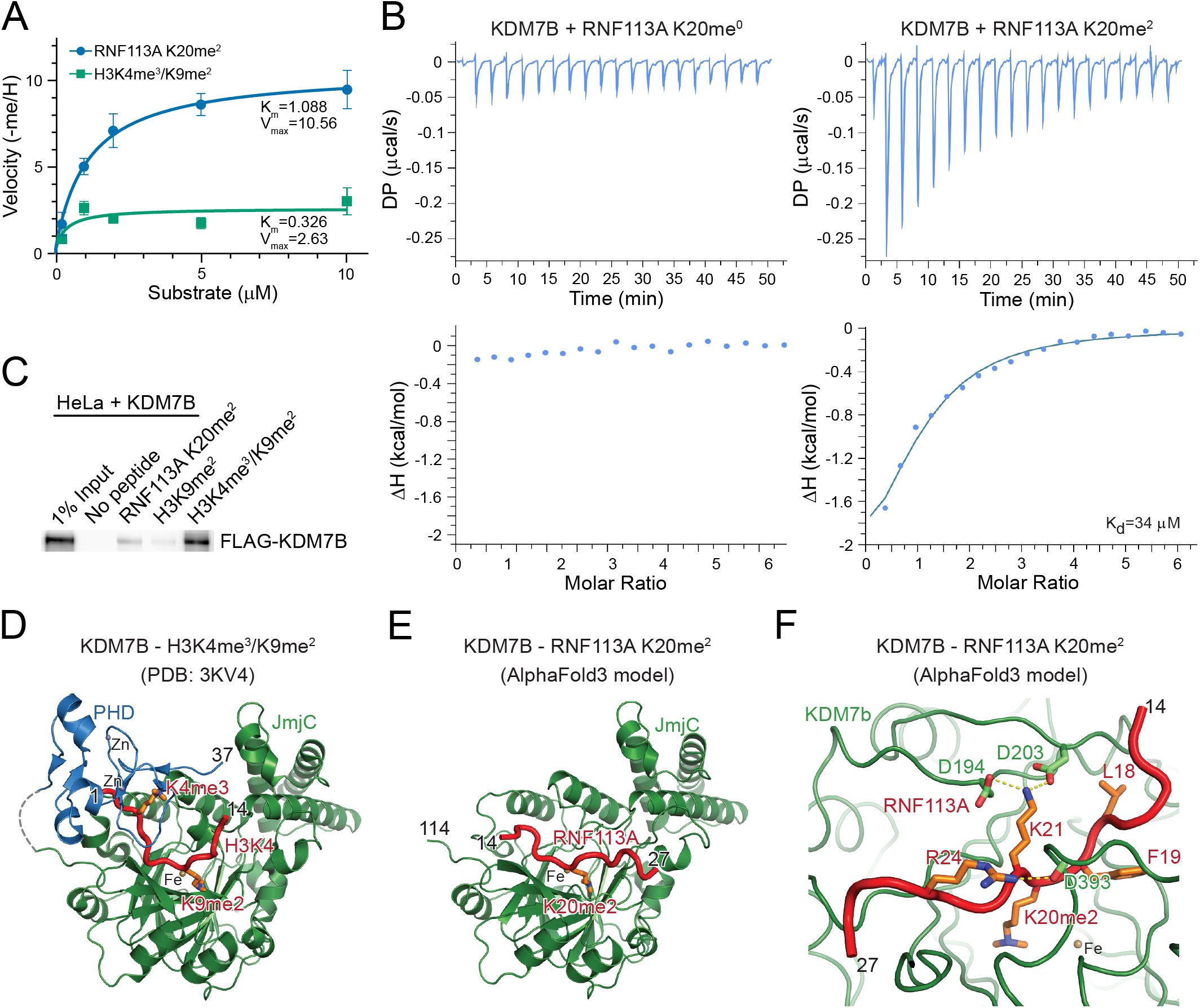
Interaction study of KDM7B-RNF113A. **A**, KDM7B enzymatic kinetics determining the Km (Michaelis-Menten constant) and Vmax (maximum enzyme velocity) of RNF113A and Histone H3 substrates according to different RNF113A or Histone H3 concentrations. Data are represented as nonlinear regression with mean ± SD of three technical replicates. **B**, ITC measurement of the interaction affinity between KDM7B (38-494) and RNF113A K20me^0^ (left panel) and RNF113A K20me^2^ (right panel) peptides. **C**, Immunodetection with indicated antibodies of peptide pulldowns using HeLa cell extracts ectopically expressing KDM7B and RNF113A K20me^2^, H3K9me^2^ and H3K4me^3^K9me^2^ peptides as bait. **D**, Crystal structure of KDM7B bound to a H3K4me^3^/K9me^2^ peptide (PDB: 3KV4). The KDM7B Jmj domain is shown in green and PHD domain in blue. Histone H3 is highlighted in red, with K4me} and K9me} shown as sticks. **E**, Alphafold3 predicted structure of the complex between KDM7B (114-482) and RNF113A-K20me^2^ (12-30). The KDM7B Jmj domain is shown in green. Only residues 14-27 of RNF113A are shown in red, with K20me^2^ shown as sticks. **F**, Details of the Alphafold3 modelled interaction between the KDM7B Jmj domain and the RNF113A-K20me^2^ peptide. RNF113A residues form several side chain interactions with KDM7B. Main chain contacts of K21, P22 and G23 with KDM7B are not shown for clarity.

### KDM7B demethylates RNF113A in cells and regulates its ligase activity

To test the capacity of KDM7B to demethylate RNF113A in cells, we expressed different combinations of tagged KDM7B, SMYD3 and RNF113A in 293T cells to monitor any potential variations in RNF113A methylation levels. Ectopic RNF113A was immunoprecipitated and its modification analyzed using a specific RNF113A K20me^3^ antibody [6]. A strong methylation signal was detected upon SMYD3 overexpression, which was significantly lower with KDM7B overexpression alone (Figure 3A). Remarkably, we observed low RNF113A methylation levels when both SMYD3 and KDM7B were expressed. This suggests that KDM7B efficiently competes with SMYD3 to impede RNF113A methylation. In addition, we noticed a clear KDM7B and RNF113A co-immunoprecipitation only in presence of SMYD3, when KDM7B actively demethylates RNF113A. Next, we tested if the effect of KDM7B was observable at physiological levels. We repeated the experiment with endogenous proteins, by generating a stable HeLa cell line with doxycycline-inducible expression of a shRNA against KDM7B. Total endogenous RNF113A was enriched from cells repressed or not for KDM7B, and the level of methylated RNF113A was analyzed using the specific RNF113A K20me^3^ antibody (Figure 3B). We observed a clear increase of RNF113A methylation following KDM7B repression, indicating that KDM7B actively demethylates RNF113A under physiological conditions. Additionally, we observed similar results when transfecting HeLa cells with specific siRNA against KDM7B (Supplementary Figure 3A).

**Figure 3:**
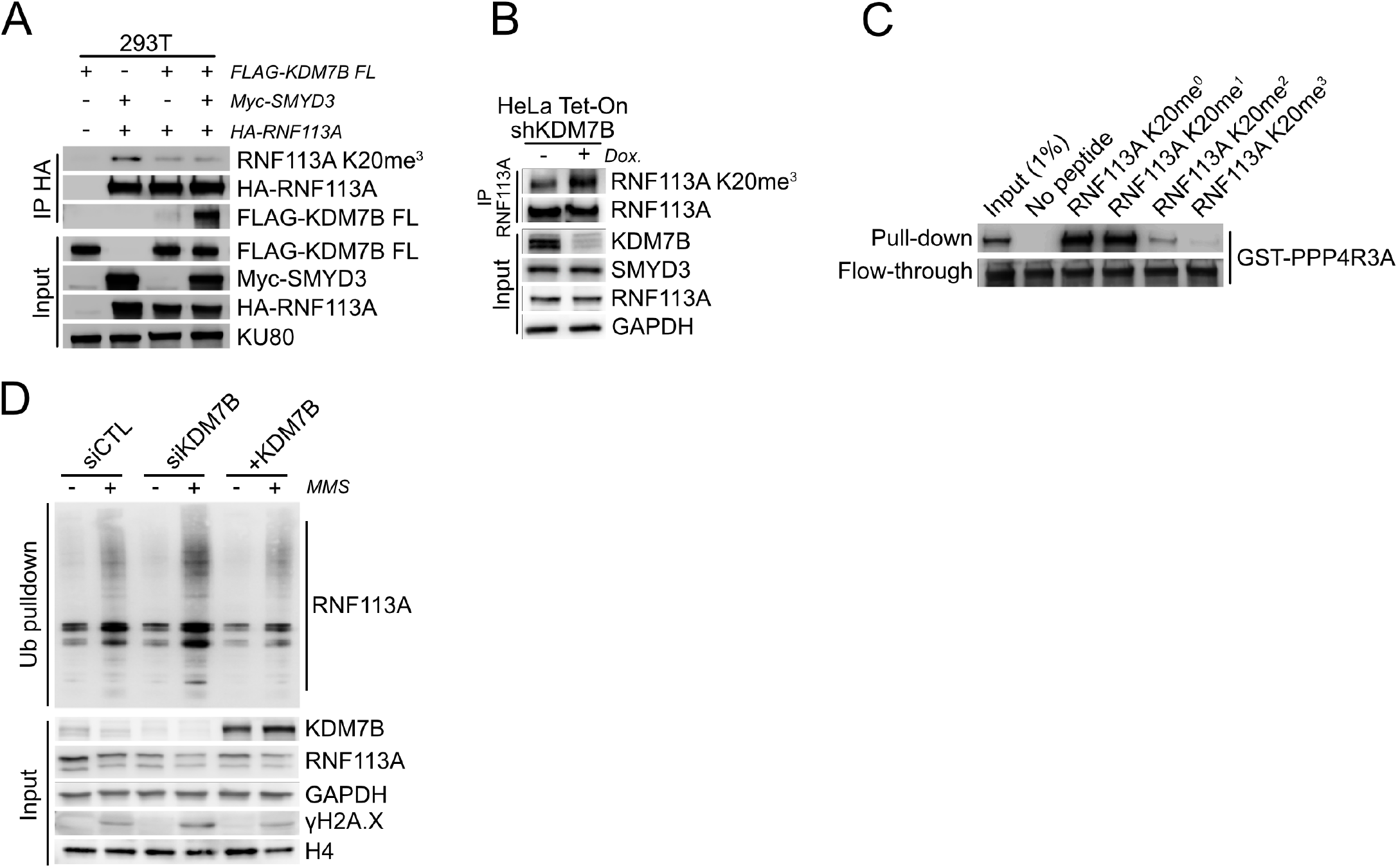
KDM7B demethylates RNF113A in cells and modulates its E3 ligase activity. **A**, Immunodetection with indicated antibodies of RNF113A demethylation by KDM7B in 293T cells after ectopic expression of HA-RNF113A, Myc-SMYD3 and Flag-KDM7B. KU80 was used as a loading control. **B**, Immunodetection with indicated antibodies of endogenous RNF113A methylation levels after immunoprecipitation of total RNF113A in HeLa cells stably engineered with an inducible shRNA targeting KDM7B upon doxycycline treatment. GAPDH was used as a loading control. **C**, Immunodetection with indicated antibodies of peptide pulldowns with recombinant GST-PPP4R3a using different states of methylated RNF113A peptides as bait. **D**, Immunodetection with indicated antibodies after TUBE pulldowns using HeLa cell extracts following repression or ectopic expression of KDM7B, in absence or presence of MMS alkylating agent. γH2A.X is shown as a marker of DNA damage induction and GAPDH and H4 were used as loading controls.

RNF113A is specifically activated by alkylating agents, leading to the stimulation of its E3 ligase activity [5]. We previously found that the methylation of RNF113A by SMYD3 methyltransferase blocks the interaction of the PP4 phosphatase and impedes RNF113A dephosphorylation. Thus, SMYD3 methylation leads to a prolonged ubiquitination activity of RNF113A as the ligase stays in a phosphorylated-active state [6]. We hypothesized that KDM7B may counteract SMYD3 regulatory functions on RNF113A activity. First, we analyzed the binding efficacy of the regulatory subunit of PP4 (PPP4R3A) with different methylated states of RNF113A. We observed by peptide pulldowns that PPP4R3A only interacts with the unmethylated or monomethylated forms of RNF113A (Figure 3C). We confirmed this result by surface plasmon resonance (SPR) analyses, observing that PPP4R3A binds strongly to RNF113A when unmodified or mono-methylated, with dissociation constants (*K*_*d*_) of 4 and 6 μM, respectively (Supplementary Figure 3B). We could not measure any affinity of PPP4R3A with di or trimethylated RNF113A. Therefore, by competing with SMYD3 to maintain RNF113A methylation level below dimethylation, KDM7B enables the phosphatase PP4 to bind and inactivate RNF113A. We then analyzed RNF113A E3 ligase activity upon KDM7B modulation in cells, by monitoring its auto-ubiquitination levels. For this, we performed ubiquitin pulldowns followed by RNF113A immunodetection using HeLa cells ectopically expressing KDM7B or treated with either a control siRNA or a siRNA targeting KDM7B. HeLa cells were incubated with methyl methanesulfonate (MMS), a potent alkylating agent, to induce alkylation damages known to activate RNF113A [6]. We observed clear increase of RNF113A autoubiquitination after MMS treatment and the RNF113A auto-ubiquitination activity was correlated to the level of KDM7B present in cells (Figure 3D and Supplementary Figure 3C). As expected, KDM7B repression led to an increase of RNF113A activity. Conversely, KDM7B overexpression resulted in decreased levels of methylated RNF113A and a corresponding reduction in the observed auto-ubiquitination signal.

Together, our data show that KDM7B actively demethylates RNF113A in cells and is able to counteract SMYD3-dependent stimulation of RNF113A E3 ligase activity.

### KDM7B regulates RNF113A functions in DNA alkylation repair

RNF113A plays a key role in DNA alkylation response through the recruitment of the ASCC dealkylation repair complex [5]. This complex repairs alkylated lesions before adducts turn into DNA double-strand breaks. Importantly, we have previously reported that RNF113A methylation by SMYD3 contributes to cell resistance to alkylating agents by enhancing DNA dealkylation repair efficacy [6]. We hypothesized that KDM7B counter-regulation of RNF113A activity may increase DNA damage and cancer cell sensitivity to alkylating treatment. We took advantage of the RNF113A-dependent ASCC complex capacity to form nuclear speckle bodies upon alkylation [5]. We generated a stable U2OS Tet-On shKDM7B cell line to efficiently monitor ASCC localization through immunofluorescence against the ASCC2 subunit, upon KDM7B repression and induction of alkylation damage by MMS exposure. We observed significantly more positive cells with foci after MMS treatment when KDM7B was repressed (25.2% vs 47.6% of cells with ≥10 foci), confirming that KDM7B is capable of downregulating RNF113A function in alkylation DNA damage response (Figure 4A-B and Supplementary Figure 4A). An impaired recruitment of the ASCC complex induced by KDM7B’s control over RNF113A is expected to increase overall DNA damage and cell sensitivity to alkylating agents, as cells fail to repair alkylation adducts before DNA replication. To test this hypothesis, we performed neutral comet assays to assess DNA integrity by quantifying olive moments (representative of the head-to-tail intensity ratio of the comet). Upon MMS treatment, we observed increased levels of DNA damage under all conditions, likely resulting from replication stress-induced double-strand breaks. However, upon a recovery period allowing the cells to repair DNA alkylation damage prior to entering their replication phase, we noticed that HeLa cells lacking KDM7B were recovering more efficiently than control HeLa cells, while DNA from cells overexpressing KDM7B was more damaged (Figure 4C-D). In parallel, we used an alternative approach to confirm KDM7B functions in HeLa cells tolerance to DNA alkylation damage. HeLa cells treated with increasing doses of MMS and depleted for KDM7B by siRNA were less sensitive to alkylation treatment compared to cells treated with a control siRNA (Figure 4E).

**Figure 4:**
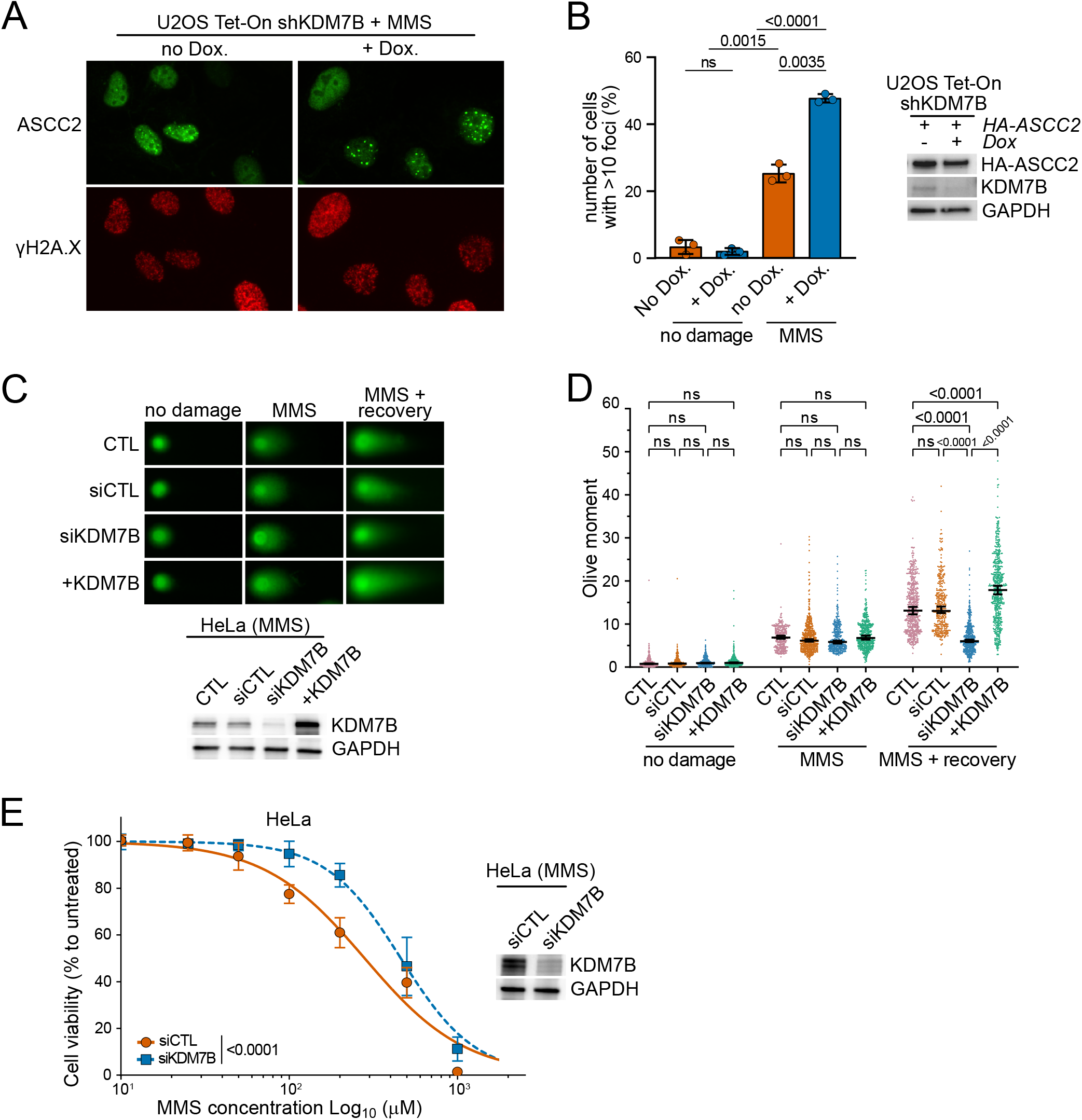
KDM7B modulates RNF113A methylation dynamics and its function in DNA dealkylation repair. **A**, Representative images of MMS-induced ASCC2 foci formation in U2OS cells stably engineered with an inducible shRNA targeting KDM7B upon doxycycline treatment, in presence of MMS alkylating agent. Foci were monitored by immunofluorescent staining of ASCC2 (upper panel) and the DNA damage marker γH2A.X (lower panel). **B**, Left panel, quantification of ASCC2 foci formation from (A). A minimum of 100 cells were analyzed for each experimental condition. *P*-values were calculated by a Dunnett’s T3 multiple comparisons test, and error bars represent mean ± SD. Right panel, immunodetection with indicated antibodies of U2OS cells expressing or not KDM7B. GAPDH was used as a loading control. **C**, Upper panel, neutral comet assays depicting HeLa cells DNA damage following exposure to MMS alkylating agent with representative examples of comet tails. Lower panel, immunoblot with indicated antibodies of cells used in the assay. GAPDH was used as a loading control. **D**, Quantification of olive moments for a minimum of 300 comets from two independent experiments for each condition. *P*-values were calculated by two-way ANOVA with the Tukey test for multiple comparisons. Data are represented as median with 95% CI. **E**, HeLa cell viability assays after treatment with different concentrations of MMS. Cells were transfected with either siRNA control or siRNA targeting KDM7B. The percentage of viable cells under each condition was normalized to vehicle-treated (control) cells. Each condition represents the mean of five technical replicates from three independent experiments. *P*-value was calculated by two-way ANOVA. Data are represented as nonlinear regression with mean ± SD. Right, immunoblot with indicated antibodies of cells used in the assay.

Our results suggest that RNF113A-dependent ASCC dealkylation response is impacted by RNF113A methylation dynamics regulated by KDM7B. KDM7B has already been linked to genome integrity through different mechanisms unrelated to dealkylation repair [24,27,28]. Thus, we decided to challenge the specificity of our observations by repeating similar comet assays upon an alternative DNA damage treatment. Remarkably, we observed that HeLa cells treated with doxorubicin, a DNA intercalating genotoxic agent, show no significant differences in DNA damage related to KDM7B expression levels (Supplementary Figure 4B-C).

Altogether, our findings demonstrate that KDM7B specifically controls RNF113A methylation level and prevents the recruitment of the DNA dealkylation repair machinery upon alkylation damage. As a result, the presence of KDM7B improves cancer cells sensitivity to alkylating agent by impairing ASCC alkylation damage response.

### KDM7B is a good prognosis marker in SCLC

Small Cell Lung Cancer is the most aggressive form of lung cancers, with limited therapeutic options. We previously found that SMYD3-dependent RNF113A methylation can be targeted to sensitize human SCLC cells and mouse models to alkylation agents and represents an interesting opportunity to improve alkylating-based chemotherapy for SCLC patients [6]. Given the importance of KDM7B in modulating RNF113A methylation dynamics and cells sensitivity to alkylation damage, we sought to evaluate its prognostic value in SCLC. We first determined the expression levels of the different actors of the RNF113A methylation signaling pathway in a representative panel of different SCLC cell lines. We noticed that the aggressive H82 cell line highly expressed SMYD3, RNF113A and KDM7B, making it an interesting model to study the functional consequences of RNF113A methylation dynamics in chemoresistance (Figure 5A). We established a doxycycline-inducible system to repress KDM7B expression in H82 cells (Figure 5B), and found that loss of KDM7B leads to better tolerance to alkylating treatment (Figure 5C). We next used these engineered H82 cells to perform xenografts in immunocompromised mice in combination with cyclophosphamide (CP), an alkylating agent frequently used in chemotherapy (Figure 5D). We observed no significant growth variations upon KDM7B repression in untreated mice, suggesting that KDM7B alone is not impacting tumor growth in our system. However, while all tumors were responsive to cyclophosphamide, tumors with intact KDM7B expression were significantly more sensitive to alkylation-based chemotherapy (Figure 5E). Further tumors analysis by immunohistochemistry (IHC) confirmed that chemotherapy was more potent in the presence of KDM7B, with a significant increase in cell death (cleaved Caspase 3 positive cells) in control tumors compared to KDM7B-depleted tumors (Figure 5F-G). As expected, no difference in cell proliferation (phosphorylated Histone H3 positive cells) was observed. These results strongly suggest that KDM7B acts as a negative regulator of RNF113A-dependent DNA dealkylation repair and that SCLC with high levels of KDM7B may be more responsive to chemotherapy.

**Figure 5:**
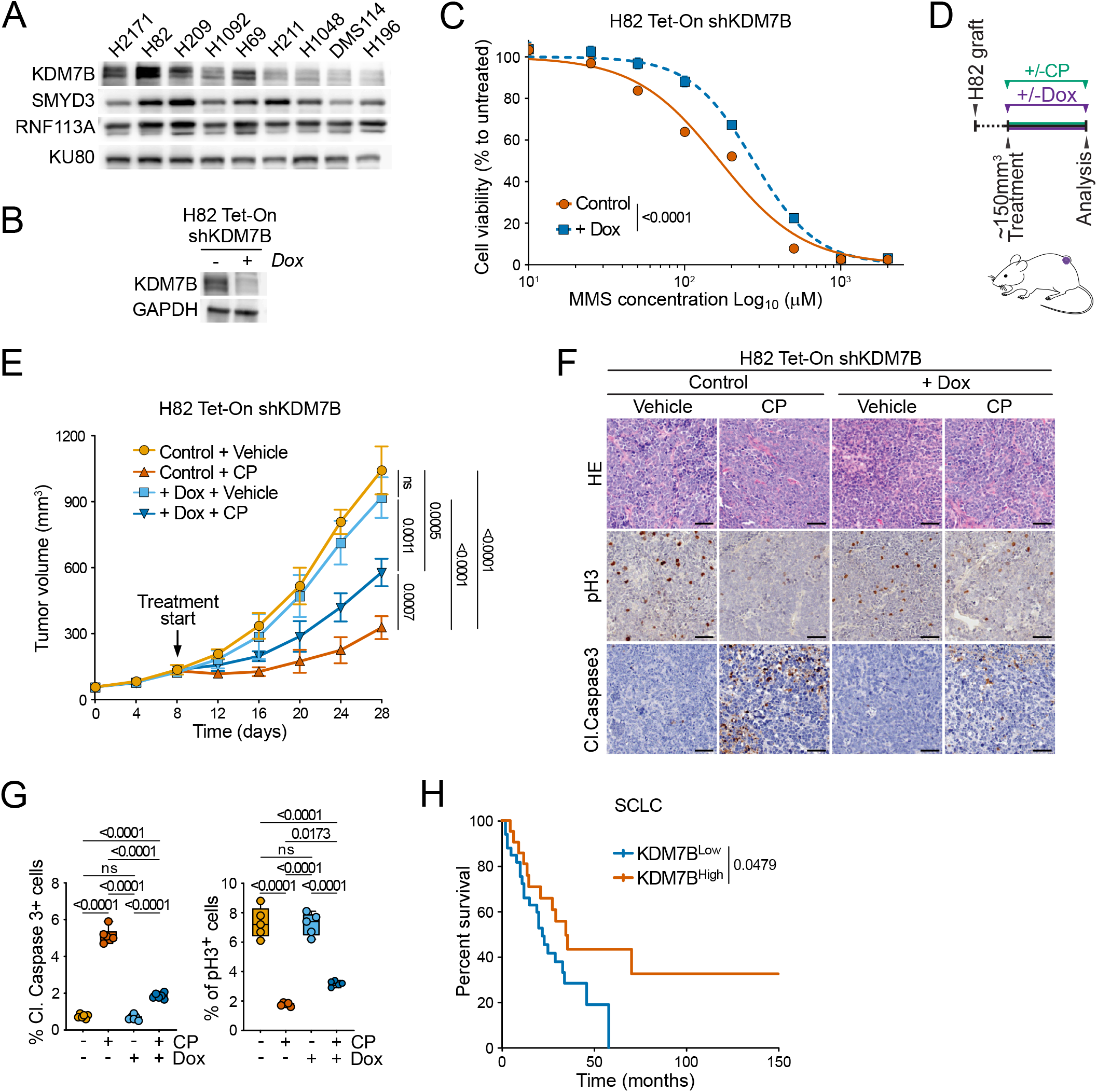
KDM7B is a good prognosis marker in SCLC. **A**, Immunoblot analysis with indicated antibodies using lysates obtained from a panel of human SCLC cell lines. KU80 was used as a loading control. **B**, immunodetection with indicated antibodies of H82 stably engineered with inducible repression of KDM7B upon doxycycline treatment. GAPDH was used as a loading control. **C**, Cell viability assays using H82 inducible shKDM7B stable cell line and different concentrations of MMS. The percentage of viable cells under each condition was normalized to vehicle-treated (control) cells. Each condition represents the mean of five technical replicates from three independent experiments. *P*-value was calculated by two-way ANOVA. Data are represented as nonlinear regression with mean ± SD. Right panel, immunoblot with indicated antibodies of cells used in the assay. **D**, Schedule protocol for H82 inducible shKDM7B stable cell line xenografts with CP or Vehicle treatment. The cells were grafted subcutaneously to immunocompromised *NOD*.*SCID-IL2Rg*^−/−^ (NSG) mice. **E**, Quantification of H82 inducible shKDM7B stable cell line xenograft tumor volume (*n* = 5 mice, for each treatment group). Animals in control groups received placebo (vehicle) treatment. *P*-values were calculated by two-way ANOVA with Tukey testing for multiple comparisons. Data are represented as mean ± SEM. **F**, Representative HE-stained sections and IHC staining for markers of cell proliferation (pH3) and apoptosis (cleaved Caspase 3) of the H82 inducible shKDM7B stable cell line xenograft tumors. Scale bars, 50 μm. **G**, Quantification of pH3, a marker of proliferation and cleaved Caspase 3, a marker of apoptosis positive cells in the xenograft tumors related to panel F. Boxes: 25th to 75th percentile, whiskers: min. to max., center line: median. *P*-values were calculated by two-tailed unpaired t test. **H**, Kaplan-Meier survival curves of SCLC patients from transcriptomic dataset [31] classified into KDM7B high and KDM7B low groups based on their relative mRNA expression levels compared to the mean of KDM7B expression. Log-rank test was performed and the corresponding *P*-value is reported.

The treatment options for SCLC remain heavily dependent on chemotherapy, which targets cancer cells by inducing DNA damage. Platinum-based therapy has been preferred to alkylating agents due to lower toxicity while maintaining similar efficacy. Unfortunately, almost all SCLC patients treated with platinum-based chemotherapy ultimately develop resistance and relapse. The development of resistance necessitates a second-line regimen, almost systematically integrating alkylating agents in SCLC. Consequently, nearly all patients undergo at least one round of alkylation-based chemotherapy, where RNF113A methylation dynamics may influence the overall therapeutic outcomes. We analyzed a SCLC transcriptomic dataset [31] and noticed that, while neither SMYD3 nor RNF113A have prognostic value for survival, SCLC patients with elevated KDM7B expression have a significantly better overall survival chance (Figure 5H and Supplementary Figure 5A). This is particularly interesting because KDM7B prognostic value seems specific to SCLC, as we did not identify significant prognostic value for KDM7B in other lung cancer subtypes such as lung adenocarcinoma (LUAD) and lung squamous cancer (LUSC), nor other cancer such as breast cancer (BRCA) (Supplementary Figure 5B).

Altogether, our results suggest that KDM7B sensitizes SCLC cancer cells to alkylating agents and is a promising determinant of SCLC patients’ survival.

### KDM7B level dictates SCLC response to alkylating chemotherapy in vivo

Our observations suggest that KDM7B influences the sensitivity of the H82 cell line to alkylation damage and may serve as a prognostic marker for survival in SCLC patients. However, while patients often receive second-line regimens that include alkylating agents, limited clinical information currently prevents us from establishing a direct correlation between KDM7B expression levels and *in vivo* tumor response to such chemotherapy. To rigorously test this hypothesis, we developed two genetically engineered mouse models (GEMMs) enabling precise modulation of KDM7B levels in SCLC tumors. We utilized previously established SCLC GEMM which harbors three conditional knockout alleles for *Rb1, Rbl2*, and *p53* (referred to as TKO). This model was originally used to demonstrate the role of the SMYD3–RNF113A pathway in mediating alkylating chemotherapy response [6]. To manipulate KDM7B expression *in vivo*, we crossed the TKO mice with CRISPR-based transcriptional modulation strains designed to either repress (CRISPRi, *H11*^*TRE-LSL-dCas9-KRAB-MeCP2;CAG-rtTA*^) or activate (CRISPRa, *H11*^*TRE-LSL-dCas9-VPR;CAG-rtTA*^) target genes in a doxycycline-inducible, guide RNA – dependent manner (Figure 6A and 6E). At eight weeks of age, TKO-CRISPRi and TKO-CRISPRa mice were infected via intratracheal aerosol delivery of a lentivirus expressing both a KDM7B-targeting sgRNA and Cre recombinase. This system allows temporal control over KDM7B repression or overexpression in SCLC tumors initiated by Cre-mediated recombination, in response to doxycycline treatment. Eighteen weeks post-infection and tumor development, mice were treated with cyclophosphamide (CP) or vehicle, in conjunction with doxycycline, to assess KDM7B’s role in modulating alkylating chemotherapy sensitivity *in vivo* (Figure 6A and 6E). All mice responded to CP with improved survival compared to vehicle-treated controls. However, mice with KDM7B repression exhibited a significantly reduced survival benefit from CP therapy – a 19.6% decrease in median survival (from 48.5 to 39 days post-treatment). In contrast, KDM7B overexpression led to a marked improvement in response to CP, with a 42.7% increase in median survival (from 48 to 68.5 days post-treatment) (Figure 6B and 6F). Importantly, consistent with our H82 xenograft experiments, modulation of KDM7B expression in the absence of CP did not significantly affect tumor growth, indicating that KDM7B does not play a major role in SCLC initiation or baseline progression. To further investigate KDM7B’s impact on tumors response to CP, we performed histological and immunohistochemical (IHC) analyses on lung tumors to assess overall burden, cell proliferation, and cell death. Of note, IHC confirmed efficient repression or overexpression of KDM7B in doxycycline-treated TKO⍰CRISPRi and TKO⍰CRISPRa tumors, respectively (Figure 6C and 6G). Upon CP treatment, tumors with repressed KDM7B expression exhibited increased tumor burden and decreased cell death compared to controls (Figure 6C–D). In contrast, overexpression of KDM7B significantly enhanced CP efficacy, leading to smaller tumor masses and elevated tumor cell death (Figure 6G–H), consistent with enhanced apoptosis in cells with impaired DNA repair of alkylation-induced damage. The modulation of KDM7B expression showed minimal effect on cell proliferation in both GEM models (Supplementary Figure 6A–B), suggesting that KDM7B’s primary influence on chemosensitivity is through regulating cell-death pathways rather than altering baseline proliferative capacity.

**Figure 6:**
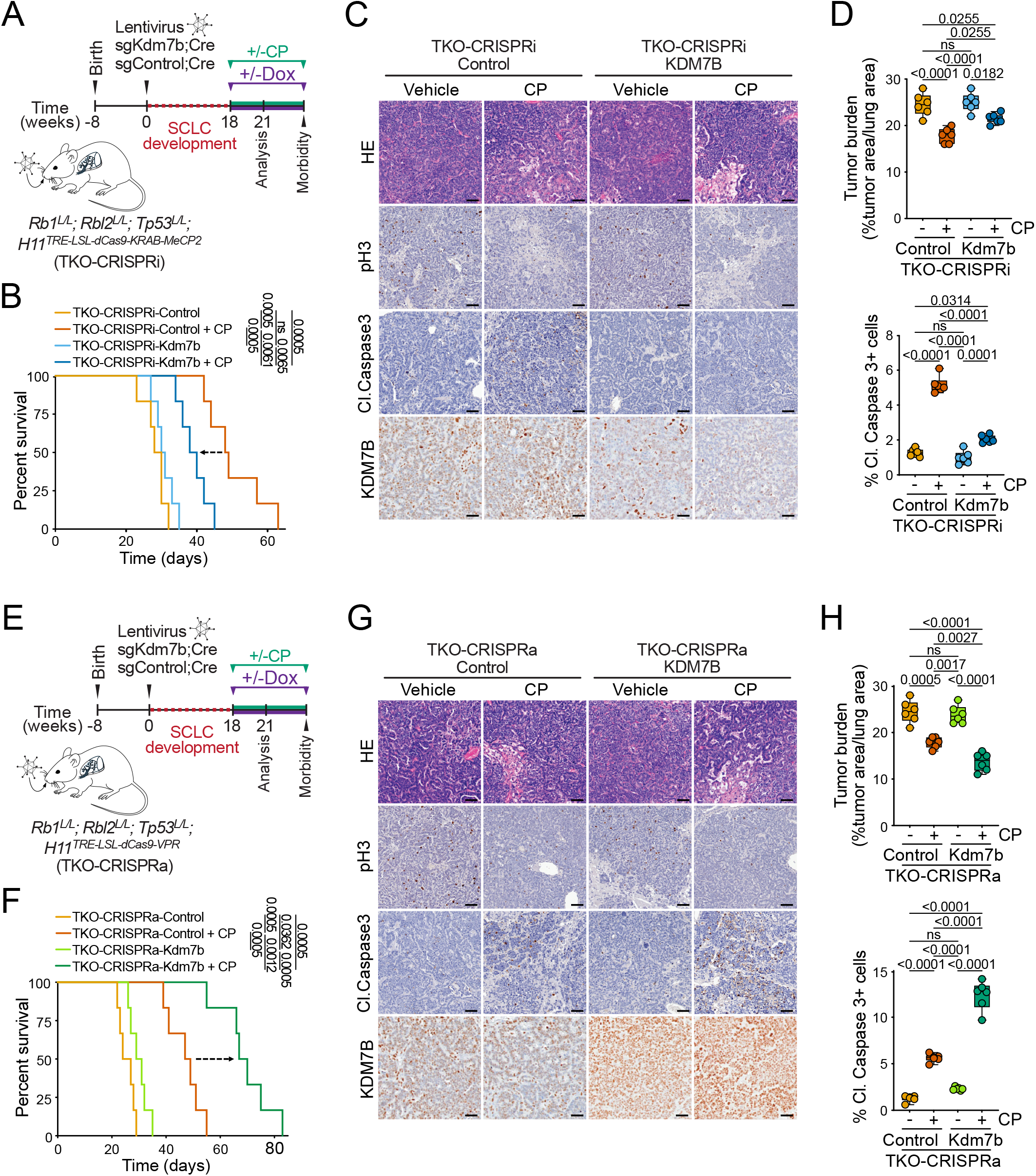
KDM7B level dictates SCLC response to alkylating chemotherapy in vivo. **A**, Schematic of treatment procedures to induce SCLC in TKO-CRISPRi mice followed by the modulation of KDM7B expression upon doxycycline treatment and the evaluation of therapeutic response to CP. TKO-CRISPRi GEMM results from the conditional deletion of *Rb1, Rbl2* and *Tp53* (TKO) and doxycycline inducible CRISPR-based repression of KDM7B upon infection with an sgKDM7B;Cre lentivirus. **B**, Kaplan– Meier survival curves of TKO-CRISPRi Control (med. survival post enrollment: 29 days, n = 5), TKO-CRISPRi Control + CP treatment (med. survival post enrollment: 48.5 days, n = 5), TKO-CRISPRi KDM7B (med. survival post enrollment: 30.5 days, n = 5) and TKO-CRISPRi KDM7B + CP treatment (med. survival post enrollment: 39 days, n = 5). *P*-values were calculated by two-way ANOVA with Tukey testing for multiple comparisons. Data are represented as mean ± SEM. **C**, Representative HE-stained sections and IHC staining for markers of cell proliferation (pH3) and apoptosis (cleaved Caspase 3) and KDM7B of TKO-CRISPRi Control and TKO-CRISPRi KDM7B tumors treated with CP or Vehicle. Scale bars, 50 μm. **D**, Quantification of tumor burden (upper panel) and cleaved Caspase 3 positive cells (lower panel), a marker of apoptosis, in the TKO-CRISPRi Control and TKO-CRISPRi KDM7B tumors treated with CP or Vehicle. Boxes: 25th to 75th percentile, whiskers: min. to max., center line: median. *P*-values were calculated by two-tailed unpaired t test. **E**, Schematic of treatment procedures to induce SCLC in TKO-CRISPRa mice followed by the modulation of KDM7B expression upon Doxycycline treatment and the evaluation of therapeutic response to CP. TKO-CRISPRa GEMM results from the conditional deletion of *Rb1, Rbl2* and *Trp53* (TKO) and doxycycline inducible CRISPR-based activation of KDM7B upon infection with an sgKDM7B;Cre. **F**, Kaplan–Meier survival curves of TKO-CRISPRa Control (med. survival post enrollment: 30 days, n = 5), TKO-CRISPRa Control + CP treatment (med. survival post enrollment: 48 days, n = 5), TKO-CRISPRa KDM7B (med. survival post enrollment: 25.5 days, n = 5) and TKO-CRISPRa KDM7B + CP treatment (med. survival post enrollment: 68.5 days, n = 5). *P*-values were calculated by two-way ANOVA with Tukey testing for multiple comparisons. Data are represented as mean ± SEM. **G**, Representative HE-stained sections and IHC staining for markers of cell proliferation (pH3) and apoptosis (cleaved Caspase 3) and KDM7B of TKO-CRISPRa Control and TKO-CRISPRa KDM7B tumors treated with CP or Vehicle. Scale bars, 50 μm. **H**, Quantification of tumor burden (upper panel) and cleaved Caspase 3 positive cells (lower panel), a marker of apoptosis, in the TKO-CRISPRa Control and TKO-CRISPRa KDM7B tumors treated with CP or Vehicle. Boxes: 25th to 75th percentile, whiskers: min. to max., center line: median. *P*-values were calculated by a two-tailed unpaired t-test.

In summary, these results identify KDM7B as a key regulator of SCLC sensitivity to alkylating chemotherapy, underscoring its potential as a predictive biomarker for determining treatment response.

## Discussion

KDM7B and RNF113A have emerged as pivotal regulators of gene expression and genomic integrity, with significant implications in cancer biology [32–34]. In this study, we identify RNF113A as a novel non-histone substrate of KDM7B and demonstrate how KDM7B-mediated demethylation of RNF113A directly regulates the alkylation damage response pathway (Figure 7). These findings broaden the functional repertoire of KDM7B, extending beyond its established roles in epigenetics and histone demethylation, and offer mechanistic insight into its unexpected involvement in the chemosensitivity of SCLC.

**Figure 7:**
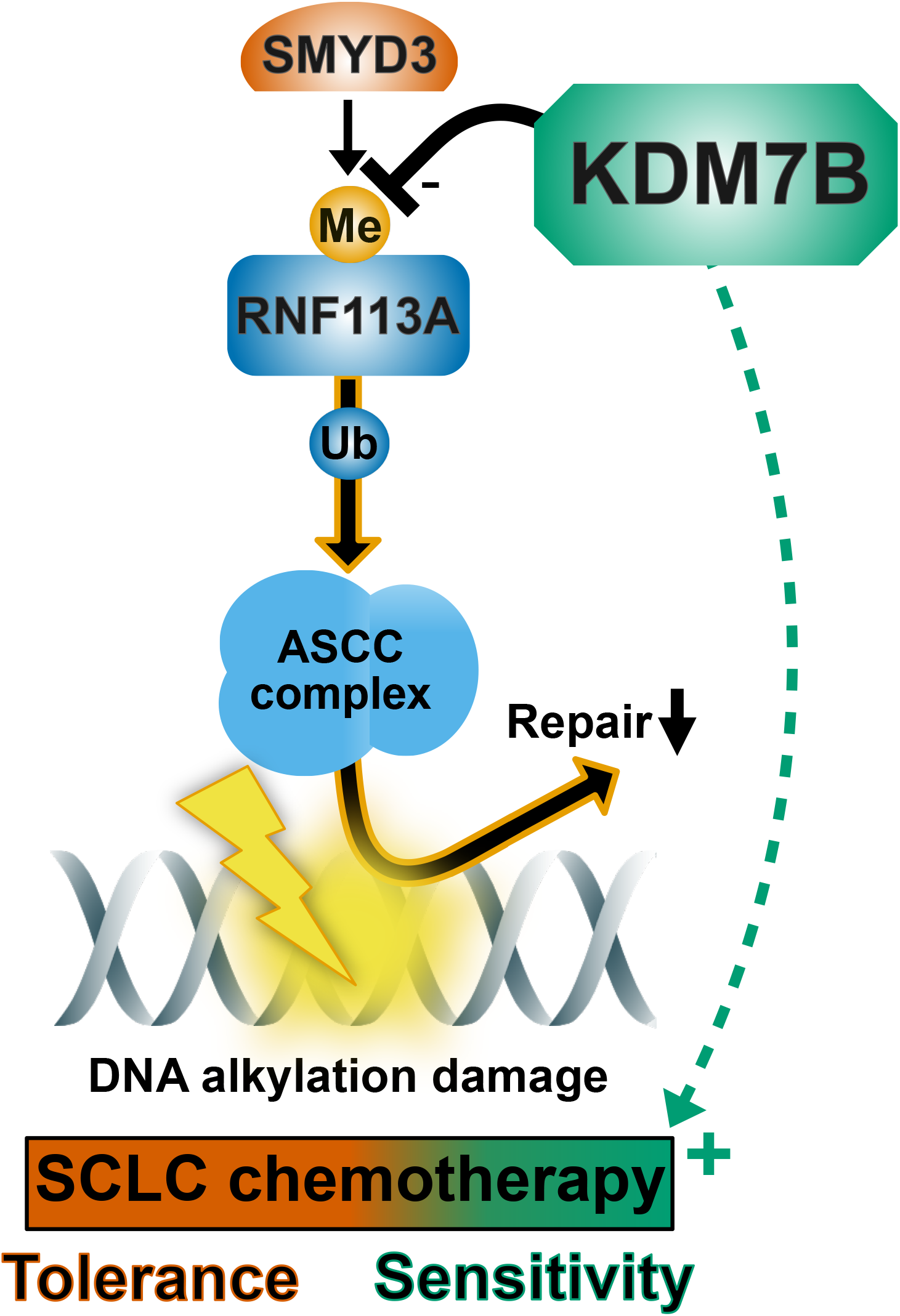
Model. Schematic representation of KDM7B control of dealkylation repair efficacy and SCLC chemosensitivity to alkylation agents through RNF113A demethylation.

Through comprehensive biochemical and functional studies, we have characterized RNF113A as a novel non-histone substrate for KDM7B. Intriguingly, our *in vitro* enzymatic assays reveal that RNF113A undergoes more efficient demethylation by KDM7B than its known histone substrate H3K9me^2^. This increased catalytic velocity is not due to a greater affinity, as indicated by isothermal titration calorimetry measurements, but rather reflects a higher turnover rate. Conversely, the demethylation of H3K9me^2^ is facilitated by KDM7B strong affinity for euchromatin environments, mediated through its PHD domain’s interaction with H3K4me^3^ [18,29]. We speculate that RNF113A may harbor an analogous stabilization motif that remains to be identified. A candidate is lysine 25 (K25), positioned at a similar spacing from methylated K20 as H3K4 is from H3K9, which could facilitate such stabilization. However, despite targeted mass spectrometry analyses, we failed to detect a K25 trimethylation event in cells, though this does not rule out the possibility of its occurrence under specific physiological conditions.

Our findings demonstrate that the methylation state of RNF113A is dependent on the combined actions of SMYD3 and KDM7B. We previously showed that SMYD3 can sequentially trimethylate RNF113A, whereas KDM7B preferentially removes mono- and dimethyl groups. Interestingly, when SMYD3 and KDM7B are co-expressed, RNF113A does not accumulate in a trimethylated state, indicating KDM7B’s competitive activity at the mono/dimethylation levels. Notably, the phosphatase PP4, essential for DNA damage signaling and RNF113A inactivation [6,35,36], fails to interact with di- or trimethylated RNF113A, highlighting the critical importance of methylation balance for RNF113A function. Thus, the interplay between SMYD3 and KDM7B delineates a crucial regulatory axis for RNF113A activity during genotoxic stress response.

KDM7B is part of a JmjC domain-containing protein subfamily that includes KDM7A and PHF2 (also known as KDM7C) [37]. Interestingly, although KDM7A shares structural similarities and exhibits similar demethylase activity on histone marks such as H3K9me^2^, H3K27me^2^ and H4K20me^1^ [18], only KDM7B was able to demethylate RNF113A in our assays. This points to a unique substrate specificity within the subfamily. Such a distinction suggests that KDM7B has evolved to target non-histone substrates relevant to stress responses, thereby expanding its biological roles beyond chromatin regulation.

KDM7B has been predominantly associated with oncogenic functions in various cancer types. It can notably promote cell migration, epithelial–mesenchymal transition (EMT), and proliferation in gastric, prostate, melanoma and breast cancers [20,22,38,39]. However, our study reveals a context-specific beneficial role of KDM7B in SCLC. Our data suggest that KDM7B alone does not impact SCLC tumor growth, but rather appears to function as a potentiator of chemotherapy by antagonizing RNF113A methylation and sensitizing cells to alkylation damage. While KDM7B appears to be a poor prognostic factor in certain cancers, there is a precedent for its beneficial role in therapeutic response. In acute promyelocytic leukemia (APL), KDM7B facilitates transcriptional reactivation of retinoic acid target genes and re-sensitizes cells to all-trans retinoic acid (ATRA), particularly in resistant cases [26]. This aligns with our findings that KDM7B is beneficial in SCLC and highlights the context-dependent functionality of KDM7B. This dual functionality of KDM7B is intriguing and may reflect the specific genotoxic stress landscape and therapeutic regimen commonly used in SCLC. It would be therefore interesting to challenge KDM7B role in other cancers commonly treated with alkylating agents.

KDM7B has previously been linked to DNA damage repair pathways, including homologous recombination (HR) in the context of double-strand break (DSB) repair and activation of the master regulator ATR under replication stress [23,28]. In those settings, KDM7B was proposed to promote chemoresistance by enhancing repair efficiency. In contrast, our data show that in the context of alkylating therapy in SCLC, KDM7B exerts a chemo-sensitizing effect. This discrepancy may reflect the specific nature of alkylating agents and their unique damage signature, which - if repaired prior to replication - does not necessarily progress to DSBs [4,40]. Our findings suggest that KDM7B acts early in the response by regulating RNF113A methylation status, potentially influencing RNA damage signaling and pre-empting more catastrophic DNA damage [34]. This highlights the significance of treatment context in understanding the role of so-called epigenetic modifiers in cancer.

Transcriptomic and proteomic analyses indicate that high KDM7B expression in SCLC patients is associated with improved prognosis, likely due to increased sensitivity to alkylation-based therapy. Although specific treatment regimens were not available in the analyzed dataset, it is highly probable, given standard clinical protocols, that most patients received cyclophosphamide or related alkylating agents as second-line therapy. This correlation suggests that KDM7B expression could serve as a predictive biomarker for response to alkylating chemotherapy. Supporting this, our *in vivo* models with inducible KDM7B modulation confirm its beneficial role in enhancing tumor sensitivity to cyclophosphamide. Given the limited efficacy and significant toxicity of current SCLC therapies, stratifying patients based on KDM7B expression may help identify those most likely to benefit from alkylating agents while avoiding unnecessary and potentially harmful treatment in non-responders.

Our *in vivo* findings also raise the possibility of therapeutically enhancing KDM7B activity during chemotherapy to boost tumor sensitivity, overcome resistance, and potentially reduce the required dose of cytotoxic agents. While direct activation of KDM7B may be challenging, precedent exists for small-molecule activators of other modifying enzymes, such as the kinase AKT and the deacetylase SIRT1 [41,42]. Alternative strategies could include stabilizing KDM7B’s interaction with RNF113A, possibly by targeting the PHD domain, or enhancing its demethylase activity through modulation of its phosphorylation level, as already described for CDK2-dependent stimulation of KDM7B activity on H3K9me^2^ [43]. Additionally, combining SMYD3 methyltransferase inhibitors with alkylation-based chemotherapy may further enhance therapeutic efficacy in tumors with high KDM7B expression by shifting RNF113A toward a demethylated and less active state, thus increasing chemosensitivity.

Although RNF113A methylation dynamics have clear implications in SCLC therapy resistance, this regulatory system may not be restricted to SCLC. RNF113A, SMYD3, and KDM7B are all highly evolutionarily conserved proteins [23,44,45], underscoring their roles in fundamental cellular processes. The evolutionary conservation of this axis also highlights its relevance beyond cancer biology. DNA bases, being highly nucleophilic, are inherently susceptible to alkylation damage. Alkylating agents are practically unavoidable, arising from both exogenous sources - such as environmental pollutants, dietary components and tobacco smoke - and endogenous processes - including oxidative stress and the activity of methyl donors like S-adenosylmethionine [40]. Therefore, cells require robust yet precisely tuned mechanisms to detect, respond to and repair these lesions, to a certain point. If left unrepaired, alkylation can compromise genome stability; however, an overactive tolerance mechanism may also result in persistence of mutations or inappropriate cell survival [46]. The KDM7B–RNF113A–SMYD3 regulatory loop appears well-positioned to maintain this delicate balance, ensuring sufficient repair while avoiding deleterious tolerance effects.

In conclusion, our study reveals a previously unrecognized beneficial role for KDM7B in SCLC, demonstrating a functional link between KDM7B and RNF113A in the response to alkylation damage. We identify a potential therapeutic window for exploiting this pathway in SCLC, in combination with chemotherapy. Further studies in larger, clinically annotated patient cohorts will be essential to validate the prognostic and predictive value of KDM7B and to develop therapeutic strategies that target RNF113A methylation dynamics.

## METHODS

### Cell culture assays

293T (RRID: CVCL_0063) and HeLa (RRID: CVCL_0030) cells were grown in DMEM (Gibco) medium supplemented with 10% FBS (Dutscher) and penicillin/streptomycin (100 U/mL Gibco). H82 (RRID: CVCL_1591), DMS-114 (RRID:CVCL_1174), H1092 (RRID:CVCL_1454), H1048 (RRID:CVCL_145), H209 (RRID:CVCL_1525), H69 (RRID:CVCL_1579), H2171 (RRID:CVCL_1536), H211 (RRID:CVCL_1529) and H196 (RRID:CVCL_1509) cells were cultured in RPMI (Gibco) medium supplemented with 10% FBS (Dutscher) and penicillin/streptomycin (100 U/mL Gibco). All cells were cultured at 37°C in a humidified incubator with 5% CO2. Cell lines were regularly checked for Mycoplasma contamination using the MycoAlert Mycoplasma Detection Kit (Lonza).

For transient expression, cells were transfected with the Mirus transfection reagent and collected 48 hours after transfection. For stable cell engineering, virus particles were produced by co-transfection of 293T cells with retroviral pMSCV FLAG/HA RNF113A WT and packaging VSVg, pΔ8.2 plasmids or with pLKO-TetOn shKDM7B using the packaging plasmids VSVg and pΔ8.2. After filtration, viruses were collected and mixed with Lenti-XTM concentrator (cat# 631232; TaKaRa Bio.). Virus/concentrator mixture was kept on ice for 30 minutes. Then virus pellet was recovered following centrifugation (1500 x g for 45 min at 4°C). Virus pellet was resuspended and used for the transduction of relevant cell lines, followed by 5 µg/mL of blasticidin or 2 µg/mL of puromycin for one week.

siRNA transfection was performed using Lipofectamine RNAi MAX (Invitrogen) according to the manufacturer’s recommendations. Final concentration of siRNA control or siRNA KDM7B (UGGGAGUGUUAGUAAUCAA, Eurogentech) was 20 nM. Cells were harvested after 72 hours of siRNA treatment.

Cells were incubated with 10 μM of the pan-demethylase IOX1 (5-carboxy-8-hydroxyquinoline; Cat. No. S7234, Selleck Chemicals) for 48 hours at 37°C prior to protein extraction and immunoblotting. To test cell survival following treatment with DNA-damaging agent (MMS: Methyl Methanesulfonate, Sigma-Aldrich), cells were cultured overnight in a 96-well plate in 100 μL media. Cells were then treated with indicated concentration of MMS for 24 hours at 37°C. In case of HeLa cells, treatment media were then replaced with standard growth media, and cell viability was assessed 72 hours later using the PrestoBlue assay (Thermo Scientific). H82 cells were kept with treatment until cell viability was assessed using the PrestoBlue assay.

### Plasmids

For recombinant protein purification, inserts coding for full length (FL) KDM7B or KDM7B 38-494 were cloned into pFL plasmid between RsrII and XbaI restriction sites with primers introducing a N-terminus Strep tag. Additionally, KDM7B 38-494 was cloned into pGex6.1 plasmid between Sma1 and Xho1 restriction sites and PP4 domain [1-118] was cloned into pet52B vector with N-terminal and C-terminal his-tag. For mammalian expression plasmids, human KDM7B, RNF113A and SMYD3 cDNAs were respectively subcloned from laboratory original plasmids into pEZY Flag, pCDNA-HA and pCDNA-MYC plasmids. For inducible repression, a specific shRNA sequence against KDM7B UTR 3’ (CTATGTTGGTTCTGACAAA) was cloned into pLKO shRNA Tet-On plasmid (RRID:Addgene_21915).

### Xenograft models

For xenograft studies, the human SCLC cell line H82 was stably transduced with pLKO shKDM7B Tet-On lentivirus. Cells were pretreated with doxycycline (100 ng/mL) for 5 days before injection, and mice were fed with 625 mg/kg doxycycline hyclate diet starting at 3 days before injection and were maintained on this diet for the remainder of the experiment. Before injection, the cells were trypsinized and dissociated into single cell suspension. The trypsin was washed with an excess growth medium, and the cells were counted. The cells were then resuspended in PBS and mixed with matrigel (1:1) at a density of 2 × 10^7^ cells per mL and kept on ice until injection. Next, 100 μL of the cell suspension was injected subcutaneously into the hind flanks of NSG mice. When tumors became palpable, they were calipered to monitor growth kinetics. For therapy studies, mice were treated as indicated with CP (40 mg/kg once per week, i.p.) in vehicle 0.9% saline. Control animals received vehicle treatment.

### Animal models

*Rb1*^*LoxP/LoxP*^, *Rbl2*^*LoxP/LoxP*^, and T*p53*^*LoxP/LoxP*^ have been described before [47–49]. Mice were in FVB/C57BL6 mix background, and we systematically used littermates as controls in all the experiments. *H11*^*LSL-dCas9-KRAB-*MeCP2;CAG-rtTA^ and *H11*^*LSL-dCas9-VPR;CAG-rtTA*^ model was generated by knock-in of the *LoxP-Stop-LoxP-dCas9-KRAB-MeCP2; CAG-rtTA* or *LoxP-Stop-LoxP-dCas9-*VP64-p65-Rta(VPR); *CAG-rtTA* cDNA-polyA cassette into *Hipp11* (*H11*) locus using methods previously described [50]. Founder animals were identified by PCR followed by sequence analysis and germline transmission confirmed by crossbreeding with FVB/C57BL6 wild-type animals. Both male and female animals were used in the experiments, and no sex differences were noted. Immunocompromised NSG mice (*NOD*.*SCID-IL2Rg*^*-/-*^, Jackson Laboratories; Strain# 005557) were utilized for xenograft studies. All NSG xenograft experiments were performed on 6 to 10-week-old female animals. In all experiments, animals were numbered, and experiments were conducted in a blinded fashion. After data collection, genotypes were revealed, and animals were assigned to groups for analysis. For treatment experiments, mice were randomized. None of the mice with the appropriate genotype were excluded from this study or used in any other experiments. All mice were co-housed with littermates (2–5 per cage) in pathogen-free facility with standard controlled temperature of 72°F, with a humidity of 30–70%, and a light cycle of 12 h on/12 h off set from 7am to 7pm and with unrestricted access to standard food and water under the supervision of veterinarians, in an AALAC-accredited animal facility at the University of Texas M.D. Anderson Cancer Center (MDACC). Mouse handling and care followed the NIH Guide for Care and Use of Laboratory Animals. All animal procedures followed the guidelines of and were approved by the MDACC Institutional Animal Care and Use Committee (IACUC protocol 00001636, PI: Mazur). Tumor size was measured using a digital caliper and tumor volume was calculated using the formula: Volume = (*width*)^2^ × *length* / 2 where *length* represents the largest tumor diameter and *width* represents the perpendicular tumor diameter. The endpoint was defined as the time at which a progressively growing tumor reached 20 mm in its longest dimension as approved by the MDACC IACUC protocol (00001636, PI: Mazur) and in no experiments was this limit exceeded.

### Small cell lung cancer mouse models

To generate tumors in the lungs of *Rb1*^LoxP/LoxP^, *Rbl2*^LoxP/LoxP^; *Tp53*^LoxP/LoxP^; *H11*^LSL-dCas9-KRAB-MeCP2; CAG-rtTA^ (TKO; CRISPRi) and *Rb1*^*LoxP/LoxP*^, *Rbl2*^*LoxP/LoxP*^; *Tp53*^*LoxP/LoxP*^; *H11*^*LSL-dCas9-VPR; CAG-rtTA*^ (TKO; CRISPRa) we used Lentivirus expressing Cre and sgKdm7b or sgControl as previously described [51]. Briefly, 8-week-old mice were anesthetized by continuous gaseous infusion of 2% isoflurane for at least 10 min using a veterinary anesthesia system. Lentivirus was delivered to the lungs by intratracheal intubation. Prior to administration, virus was precipitated with calcium phosphate to improve the delivery of Cre by increasing the efficiency of viral infection of the lung epithelium. Mice were treated with one dose of 5 × 10^5^ PFU of Lentivirus-sgKdm7b;Cre and Lentivirus-sgControl;Cre. Mice were analyzed for tumor formation and progression at the indicated timepoints after viral infection. Mice were fed a standard or 625 mg/kg Doxycycline hyclate diet starting at 18 weeks after Lentivirus infection and were maintained on this diet for the remainder of the experiment. For therapy studies, mice were treated as indicated with CP (40 mg/kg once per week, i.p.) in vehicle 0.9% saline. Control animals received vehicle treatment.

### Histology and immunohistochemistry

Tissue specimens were fixed in 4% buffered formalin for 24 hours and stored in 70% ethanol until paraffin embedding. Hematoxylin and eosin (HE) staining and immunostainings were performed on 3 μm thick tissue sections. Immunohistochemistry (IHC) was performed on formalin-fixed, paraffin-embedded tissue (FFPE) sections using a biotin-avidin HRP conjugate method (Vectastain ABC-HRP kit, #PK4000) as described before^102^. The following antibodies were used (at the indicated dilutions): cleaved Caspase 3 (RRID: AB_2070042, CST, 1:100), phospho Histone 3 (RRID: AB_331535, CST, 1:1000) and KDM7B (RRID: AB_1264338, Bethyl, 1:200). Sections were developed with DAB and counterstained with hematoxylin. Pictures were taken using a PreciPoint M8 microscope equipped with the PointView software and quantified using Image J software (v1.53k, RRID:SCR_003070) and QuPath (v0.5.1, RRID:SCR_018257).

### Expression and purification of recombinant proteins

Recombinant proteins were purified from Escherichia coli BL21 and BL21Star (DE3, Invitrogen). For GST-based purification, cells were resuspended in lysis buffer (50 mM Tris pH 7.5, 150 mM NaCl, 0.05% NP-40, 0.25 mg/mL lysozyme, 0.5 mM PMSF and protease inhibitors) and were lysed by sonication. The soluble fraction was obtained by centrifugation. GST-tagged proteins were purified using Glutathione Sepharose 4B beads (GE Healthcare) and eluted with 10 mM reduced L-glutathione (Sigma-Aldrich). For His-PP4 domain [1-118], bacteria cell pellets were lysed in a buffer containing 20mM Tris pH 8, 100 mM NaCl and 2% glycerol. Lysate was clarified by centrifugation and the protein was first purified on Ni2+-Chelating Sepharose (Cytiva) and further purified by a gel filtration on Superdex 75 Increase (Cytiva).

Recombinant Proteins were alternatively produced in Hi5 insect cells. Briefly, KDM7B-FL and KDM7B (38-494) were cloned into pFL vector as Strep-tag fusion [52], and expressed using baculovirus system in Hi5 insect cells using Express five medium (Thermo). Insect cell pellets were resuspended in a lysis buffer containing 50mM HEPES pH 7.5, 300mM NaCl, 0.2mM TCEP, 0.04%, Triton X-100, 5% glycerol and Complete EDTA free protease inhibitor (Roche), and lysed by sonication. Following centrifugation for 30 minutes at 45000g at 4°C, the supernatant was applied onto a Strep-Tactin XT high capacity resin (IBA Lifesciences). KDM7B was eluted with 50mM D-Biotine (Euromedex) and further purified by gel filtration on Superdex 200 Increase column (Cytiva).

### Peptides

RNF113A peptides were purchased from Covalab: Biotin—Ahx—DQVCTFLF Kme^0^ KPGRKG—CONH2, Biotin—Ahx—DQVCTFLF Kme^1^ KPGRKG—CONH2, Biotin—Ahx—DQVCTFLF Kme^2^ KPGRKG—CONH2, Biotin—Ahx—DQVCTFLF Kme^3^ KPGRKG—CONH2. H3K9me^2^ and H3K4me^3^K9me^2^ peptides were purchased from Caslo: ARTKQTAR Kme^2^ STGGKAPRKQLAGGYK-Biotin-NH2, ART Kme^3^ QTAR Kme^2^ STGGKAPRKQLAGGYK-Biotin-NH2.

### Demethylation assays

Demethylating activity of KDM7B towards peptides (RNF113AK20me^2^, H3K9me^2^, H3K4me^3^K9me^2^) was monitored by Mass Spectrometry (MS). Briefly, 25 µL demethylase reactions (50 mM HEPES pH 7.0, 10 µM peptide, 100 µM sodium ascorbate, 10 µM 2-oxoglutarate, 25 µM Fe(II) SO4, 1 mM TCEP, 1% v/v DMSO) were performed at 37 °C for 4 hours. Demethylase activity of KDM7B enzyme towards individual peptides was performed with 1 μg of recombinant KDM7B enzyme. Peptide samples (RNF113AK20me^2^, RNF113AK20me^3^, H3K9me^2^, H3K4me^3^K9me^2^) were desalted using C18 ZipTips. For MALDI-MS analyses, peptides were mixed 1:1 with a saturated α-cyano-4-hydroxycinnamic acid (CHCA) matrix in 50% acetonitrile/0.1% TFA. One microliter of the mixture was spotted onto a MALDI plate and air-dried. Spectra were acquired on a MALDI-TOF in positive ion reflector mode. External calibration was performed using a peptide standard. Data were analyzed using DataExplore software to detect changes in peptide molecular weight. For analyses using nanoliquid chromatography coupled to electrospray-MS, peptides were sampled on a precolumn (300 μm x 5 mm PepMap C18, Thermo Scientific) and separated in a 75 μm x 250 mm C18 column (Aurora Generation 3, 1.7μm, IonOpticks) using a 20 min acetonitrile gradient. The MS and MS/MS data were acquired by Xcalibur (Thermo Fisher Scientific). The data were manually processed using Xcalibur Qual Browser (Thermo Fisher Scientific). Masses corresponding to respectively the 4+, 5+ and 6+ ions of the me0, me1 and me2 potential forms of RNF13A-K20, H3K9 and H3K4K9 peptides were extracted from MS1 scans (tolerance: 5 ppm) and their corresponding signals integrated over their complete elution windows.

Demethylating activity of KDM7B towards peptides (RNF113AK20me^2^, H3K9me^2^ and H3K4me^3^K9me^2^) was also monitored with the Succinate-Glo JmjC Demethylase/Hydroxylase Assay Kit (Cat# V7990; Promega) with minor modifications. 25 µL demethylase reactions (50 mM HEPES pH 7.0, 10 µM peptide, 100 µM sodium ascorbate, 10 µM 2-oxoglutarate, 25 µM Fe(II)SO4, 1 mM TCEP, 1% v/v DMSO) were performed at 37 °C for 1 hour. Demethylase activity of KDM7B enzyme towards individual peptides was performed with 1 μg of recombinant KDM7B enzyme ( Hi5insect cells purified KDM7B, BL21 bacteria purified GST-KDM7B). Subsequent detection of succinate was determined exactly as described by the manufacturer (Promega). Briefly, 25 µL of Detection Reagent I was added to each demethylase reaction and incubated for 1 hour, followed by the addition of 50 µL of Detection Reagent II. Luminescence was read after 10-20 min with a BMG LabTech CLARIOstar Plus Multi-mode plate reader.

### Isothermal titration calorimetry (ITC)

KDM7B (38-494) was purified using Superdex 200 Increase (Cytiva) in a buffer containing 25mM HEPES pH7.5, 100mM NaCl, 0.2mM TCEP and 5% glycerol. RNF113A (12-26) peptides were resuspended in the same buffer and KDM7B and the peptides were dialysed against this buffer overnight. ITC experiments were performed at 25 °C using a Microcal PEAQ-ITC microcalorimeter (Malvern). Experiments included one 0.5 μl injection and 19 injections of 2 μL of 1.3-1.5mM RNF113A peptides into the cell containing 50µM KDM7B. The initial data point was deleted from the data sets. Binding isotherms were fitted with a one-site binding model by nonlinear regression using MicroCal PEAQ-ITC Analysis software (Malvern).

### Surface plasmon resonance (SPR)-based biosensor analysis

The interaction of PP4-his with the different synthetic peptides was analysed by SPR using BIACORE T200. For each peptide interaction, the biosensor experiment was repeated three times over PP4-his surfaces (around 120 RU immobilized). A separate control flow cell was activated and blocked to correct for refractive index changes. The experiments were performed at 25°C using a flow rate of 15μl/min. For each peptide, solutions at four different concentrations were injected for 20 seconds over the PP4-his and control flow cells and allowed to dissociate for 10 seconds. All peptide dilutions were achieved in running buffer (100 mM NaCl, 20 mM Tris pH 8, 2% glycerol, complemented with 0.05% Tween 20 and 50 µM EDTA). After each injection, regeneration was conducted using two pulses of 10 mM NaOH and 350 mM EDTA. The dissociation constant was calculated by fitting the data with a Hill model in Prism (GraphPad). Measurements were performed in duplicate.

### Peptide pulldown

A volume of 7.5 μL of streptavidin Sepharose beads (GE Healthcare) were saturated with 10 μg of specific biotinylated peptides in peptide buffer (50 mM Tris pH 7.9, 150 mM NaCl, 0.5% NP40, 1 mM PMSF, complete protease inhibitors; Roche) for 2 hours at 4°C with rotation. Following incubation, beads were washed in the peptide buffer and incubated with either 1 mg of whole-cell extract or 1 ug of recombinant protein in peptide buffer for 3 hours at 4°C with rotation. Subsequently, beads were washed 5 times in peptide buffer. Beads were then eluted in Laemmli buffer and analyzed by immunoblotting.

### Immunoblot analysis

Proteins were separated by SDS-PAGE, transferred to PVDF membrane, and analyzed by immunoblot. The following antibodies were used: RNF113A (RRID:AB_2646617, Invitrogen; RRID:AB_1079821, Atlas/Sigma-Aldrich), KDM7B (RRID:AB_1211498, Bethyl; RRID:AB_2793316, Active Motif), FLAG (RRID:AB_259529, Sigma-Aldrich), HA (RRID:AB_10691311, CST; RRID:AB_ 2565006, Biolegend), pH2A.X (RRID:AB_2118009, CST; RRID:AB_2793161, Active Motif); H4 (RRID:AB_1147658, CST), α-tubulin (RRID:AB_1904178, CST), GAPDH (RRID:AB_1080976, GeneTex), Ku-80 (RRID:AB_2218736, CST), YAP (D8H1X) XP (RRID:AB_2650491, CST), ASCL1 (E7N9C) (RRID:AB 2943074, CST), NEUROD1 (RRID:AB_1845331, Sigma-Aldrich), POU2F3 (RRID:AB_1855585, Sigma-Aldrich), RNF113A K20me^3^ (generated by Eurogentec, speedPTM protocol), and SMYD3 (developed in house, as previously described in [53]).

### Immunoprecipitation

HA immunoprecipitation. Following cell culture and potential transfections, media was removed and cells were washed with ice-cold PBS. cells were resuspended in lysis buffer (20 mM Tris-HCl pH 7.9, 150 mM NaCl, 1 mM EGTA, 1 mM EDTA, 0.5% Triton, 1 mM PMSF, protease inhibitor) and sonicated. The cell lysate was cleared by centrifugation. The cell lysate was added to previously washed anti-HA resin (Pierce) in the same buffer. Cell lysate/resin mixture was incubated for 4 hours at 4°C with rotation. HA resin with bound proteins was washed five times in the same buffer. Proteins were eluted with Laemmli buffer and analyzed by immunoblotting using the described antibodies.

For other immunoprecipitation experiments with KDM7B repression, cells were treated with doxycycline (100ng/mL) for 48 hours or transfected with siRNA for 72 hours. Cells were collected and resuspended in lysis buffer as described before. Immunoprecipitation was performed by using anti-RNF113A antibody. Cell lysate/antibody mixture was incubated for 4 hours at 4°C with rotation. Proteins were eluted and analyzed by immunoblotting using the described antibodies.

For peptide pulldowns, 10 μL of streptavidin Sepharose beads (GE Healthcare) were saturated with 7.5 μg of specific biotinylated peptides in peptide buffer (50 mM Tris pH 8, 150 mM NaCl, 0.5% NP40, 0.5 mM DTT, 10% glycerol and complete protease inhibitors; Roche) for 2 hours at 4°C with rotation. Next, beads were washed in the peptide buffer and incubated with 1 μg of recombinant proteins in peptide buffer for 4 hours at 4°C with rotation. Beads were then washed 3 times in peptide buffer and subsequently eluted in Laemmli buffer and analyzed by immunoblotting.

### TUBE pulldown

Alkylation damage was induced with MMS treatment for 4 hours. Cells were harvested and lysed in TUBE lysis buffer (50 mM Tris-HCl pH 7.9, 1 mM EGTA, 1 mM EDTA, 1% Triton X-100, 0.27 M Sucrose, 0.2 mM PMSF, 100 mM iodoacetamide, protease inhibitors) for 1 hour at 4°C. Prior to this step, a fraction of the cell pellet was used to confirm DNA damage by pH2A.X immunoblotting. Whole-cell lysate was then added to TUBE2 beads (Recombinant Human Ubiquitin 1 Tandem UBA Agarose, R&D Systems). Samples were incubated overnight at 4°C. Beads were washed 3 times in high salt TAP buffer (50 mmol/L Tris-HCl pH 7.9, 300 mM KCl, 5 mM MgCl2, 0.2 mM EDTA, 0.1% NP-40, 10% glycerol, 2 mM β-mercaptoethanol, 0.2 mM PMSF) and twice in low-salt TAP buffer (50 mM Tris-HCl pH 7.9, 5 mM MgCl2, 0.2 mM EDTA, 0.1% NP-40, 10% glycerol, 2 mM β-mercaptoethanol, 0.2 mM PMSF). Proteins were eluted in Laemmli buffer, and TUBE pulldowns were analyzed by western blot.

### Immunofluorescence microscopy

For MMS-induced foci analysis, U2OS cells expressing a doxycycline-inducible shKDM7B cells were transduced with HA-ASCC2 and shRNA expression was induced with doxycycline for 72 hours. Cells were treated with 500 μM MMS in the complete medium at 37°C for 6 hours, washed with cold PBS, and then permeabilized with PBS containing 0.2% Triton X-100 and protease inhibitor cocktail (Pierce) for 20 minutes. After an additional washing with cold PBS, cells were fixed with 3.2% paraformaldehyde in PBS. Cells were then washed extensively with IF Wash Buffer (PBS, 0.5% NP-40, and 0.02% NaN3), then blocked with IF Blocking Buffer (IF Wash Buffer with 10% FBS) for at least 30 minutes. Primary antibodies (anti-HA, anti-pH2AX) were diluted in IF Blocking Buffer overnight at 4°C. After staining with secondary antibodies (conjugated with Alexa Fluor 488; RRID:AB_2534069, Invitrogen; conjugated with Alexa Fluor 594; RRID:AB_2534095, Invitrogen), samples were mounted using the Prolong Gold mounting medium (Invitrogen). Epifluorescence microscopy was performed using an Olympus fluorescence microscope (BX-53) with an UPlanS-Apo 100×/1.4 oil immersion lense and cellSens Dimension software. For foci quantification, at least 100 cells were counted in three biological replicates.

### Neutral comet assay

HeLa cells were transfected with either siRNA siKDM7B or siCTL or KDM7B plasmid for ectopic expression 48 hours prior to treatment with either 1 mM MMS for 1 hour or 5 µM doxorubicin for 2 hours, as indicated. Media were replaced with fresh media and cells were further incubated for 2 hours when indicated. Cells were then washed once with PBS and trypsinized. Cells were resuspended in ice-cold PBS at a concentration of 1 × 10^5^ cells/mL. A neutral comet assay was then performed using the CometAssay (Trevigen) kit, according to the manufacturer’s protocol. Briefly, cells were mixed with low melting point agarose at a ratio of 1:10, and then 80 μL of this mixture was spread onto a comet slide and incubated at 4°C for 30 minutes. Slides were immersed in ice-cold lysis buffer (Trevigen) for 1 hour at 4°C, then in 1X TBE buffer (0.1M Tris Base, 0.1M boric acid, 2.5 mM EDTA) for 30 minutes at 4°C. After lysis, cells were electrophoresed in 1X TBE buffer at 20 V for 30 minutes at 4°C. Slides were rinsed with distilled water and incubated with DNA precipitation solution (1M ammonium acetate in 95% ethanol) for 30 minutes at room temperature. Slides were then incubated with 70% ethanol for 30 minutes and dried overnight at room temperature in the dark. DNA staining was done using 1X SYBR Gold (Thermo Fisher) at room temperature for 30 minutes. After drying the slides, images were acquired with an epifluorescence microscope (Zeiss; 10X; AxioVision control software, RRID:SCR_002677). A minimum of 300 comets were scored for each condition using the OpenComet plugin in ImageJ (RRID:SCR_003070).

### Statistical analysis

Graphpad Prism software was used to perform the statistical analyses using the statistical methods presented in the respective figure legends.

### Bioinformatics analysis

Transcriptomic data were obtained from EGA study EGAS00001000925, EGA dataset EGAD00001001244: 118 paired .fastq files corresponding to 59 patients. Clinical data were obtained from [31] (41586_2015_BFnature14664_MOESM72_ESM.xlsx file). The sequenced reads from the raw sequence (.fastq files) were aligned on the UCSC hg38 Homo_sapiens genome using the STAR software (2.7.1a) [54] to produce bam files. The aligned reads (.bam files) were counted using HTSeq framework (0.11.2) [55], with options: -t exon -f bam -r pos –stranded=no -m intersection-strict –nonunique none. The normalizations were performed using the R version 4.4.2 (2024-10-31) and DESeq2 (1.46.0) [56,57] and SARTools (1.8.1) [58] packages.

## DATA AVAILABILTY

All gene-expression data utilized in the study are publicly available (from EGA study EGAS00001000925, EGA dataset EGAD00001001244). This study did not generate any unpublished code, software, or algorithm. Any additional information required to reanalyze the data reported in this paper is available upon request from the corresponding author.

Unique biological materials created for this study will be available from the corresponding author upon request.

## ACKNOWLEDGMENT

IBS acknowledges integration into the Interdisciplinary Research Institute of Grenoble (IRIG, CEA). This work used the platforms of the Grenoble Instruct-ERIC Center (ISBG; UAR 3518 CNRS-CEA-UGA-EMBL) within the Grenoble Partnership for Structural Biology (PSB), supported by FRISBI (ANR-10-INBS-0005-02) and GRAL, financed within the University Grenoble Alpes graduate school (Ecoles Universitaires de Recherche) CBH-EUR-GS (ANR-17-EURE-0003). We thank Caroline Mas for assistance with ITC. MS-based experiments were partially supported by Agence Nationale de la Recherche under projects ProFI (Proteomics French Infrastructure, ANR-10-INBS-08) and GRAL, a program from the Chemistry Biology Health (CBH) Graduate School of University Grenoble Alpes (ANR-17-EURE-0003). IAB acknowledges the EpiMed core facility (http://epimed.univ-grenoble-alpes.fr) for their support and assistance in the bioinformatics analysis. Most of the computations presented in this paper were performed using the GRICAD infrastructure (https://gricad.univ-grenoble-alpes.fr), which is supported by Grenoble research communities.

S.H. was supported by the NIH (K99 CA255936). N.M.F. was supported by the American Cancer Society postdoctoral fellowship. X.L. by the CPRIT Training Award (RP210028). F. Lan was supported in parts by NSFC 32525001 and 32430022. N. Mosammaparast was supported in parts by the Siteman Cancer Center and the Barnard Foundation. P.K. Mazur was supported in part by grants from the NIH (R01 CA272844, R01 CA278940, R01 CA236949, R01 CA266280, R01 CA272843), DoD PRCRP Career Development Award (CA181486), CPRIT IIRA (RP220391) and CPRIT Scholar in Cancer Research (RR160078). N. Reynoird was supported in parts by the INCa (PLBIO19-021), the Ligue contre le cancer (comité Savoie), the APMC (Agir Pour les Maladies Chroniques), and the Fondation ARC (PJA 20181207702).

## AUTHOR CONTRIBUTIONS

**T. Ahmad:** Investigation, visualization, writing – original draft, writing – review & editing. **X. Yang**: Investigation and visualisation. **AE. Foucher:** Investigation, visualization, writing – review & editing. **L. Ren:** Investigation, visualization, writing – review & editing. **N. Tsao:** Investigation, visualization, writing – review & editing. **NM. Flores:** Investigation and visualisation. **J. Wan:** Investigation and visualization. **E. Dubiez:** Investigation and visualisation. **L. Belmudes:** Investigation and visualization **MC. Bhowmik:** Investigation. **J. Vayr:** Investigation. **SC. Hausmann**: Investigation and visualization. **F. Chuffart:** Investigation. **X. Lu**: Investigation and visualization. **S. Blanchet:** Investigation. **T. Chasan**: Investigation and visualization. **F. Boussouar:** writing – review & editing. **Y. Coute:** Supervision, writing – review & editing. **N. Mosammaparast:** supervision, funding acquisition, writing – review & editing. **F. Lan:** Supervision, funding acquisition, writing – review & editing. **J. Kadlec:** Supervision, funding acquisition, writing – review & editing. **P.K. Mazur:** Supervision, funding acquisition, writing – review & editing. **N. Reynoird:** Conceptualization, supervision, project administration, funding acquisition, visualization, writing – original draft, writing – review & editing.

