## Supplementary figures and images for "KDM7B-mediated demethylation of RNF113A regulates small cell lung cancer sensitivity to alkylation damage"

### Supplementary Figure 1

#### SUPPLEMENTARY FIGURE 1

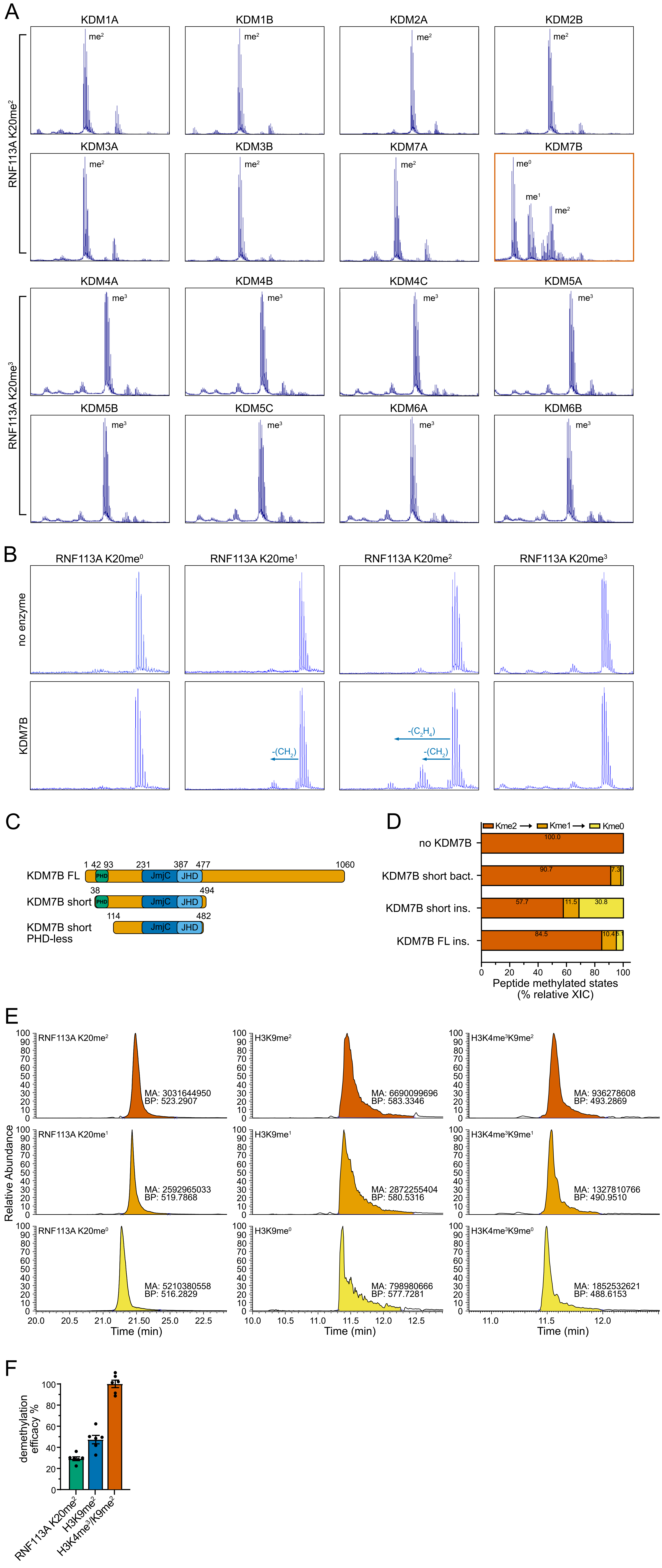

### Supplementary Figure 2

#### SUPPLEMENTARY FIGURE 2

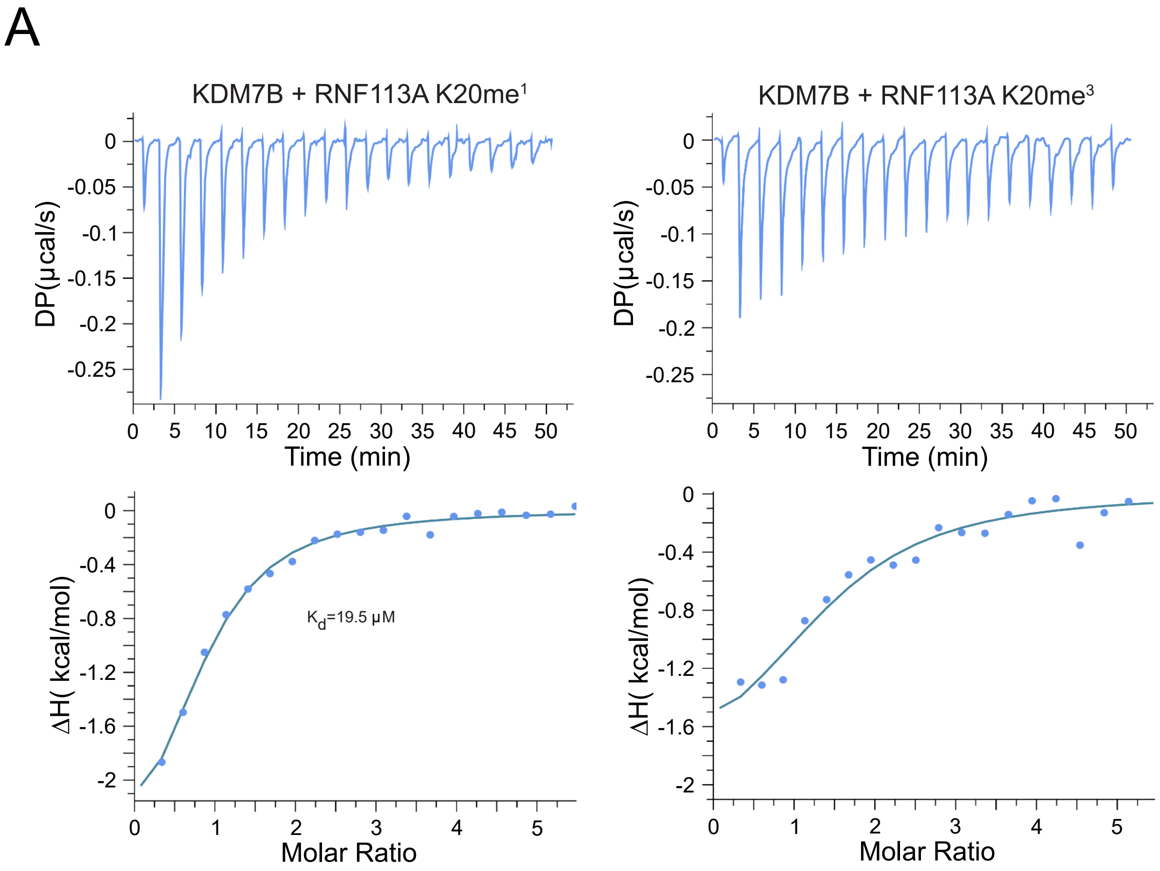


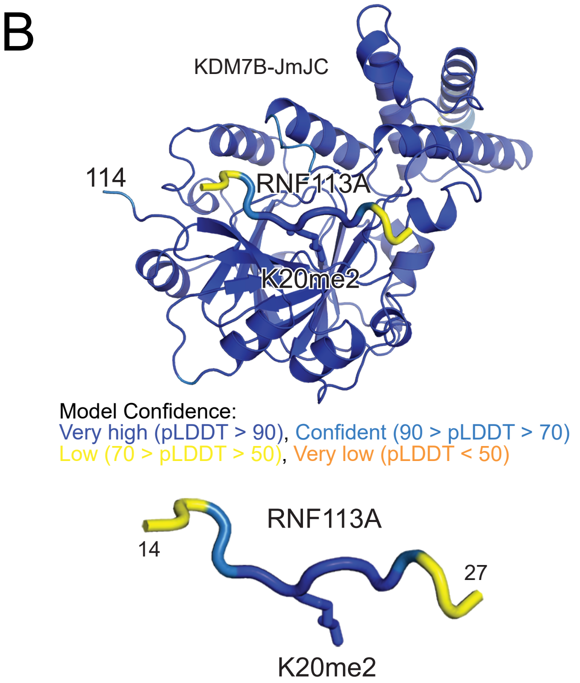


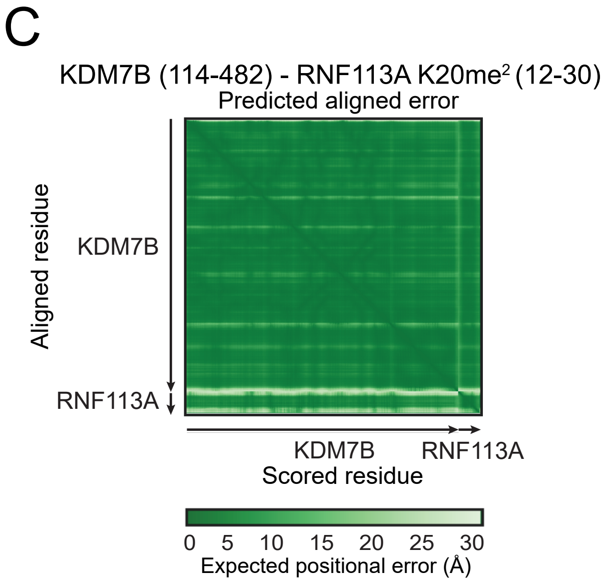

### Supplementary Figure 3

#### SUPPLEMENTARY FIGURE 3

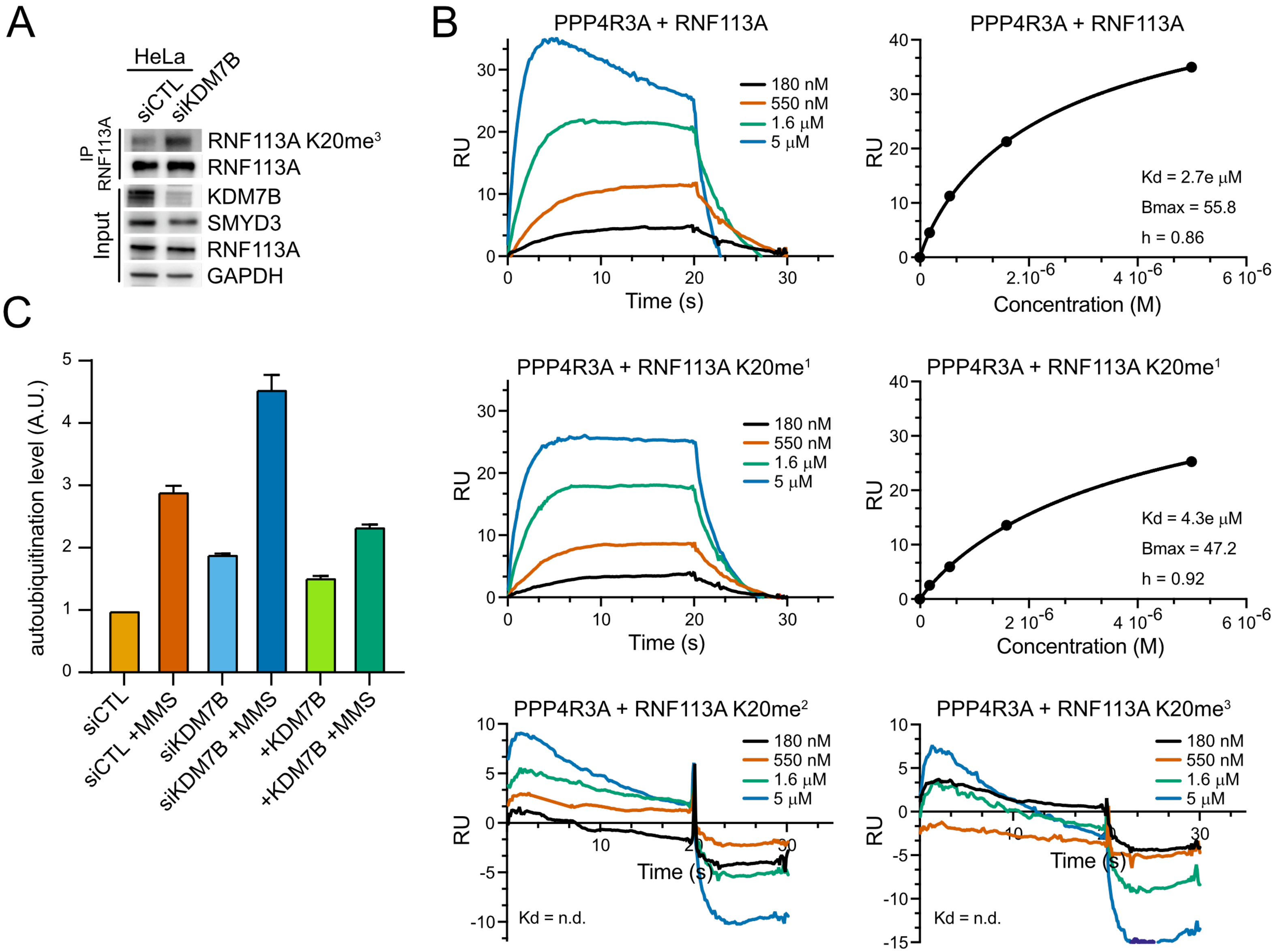

### Supplementary Figure 4

#### SUPPLEMENTARY FIGURE 4

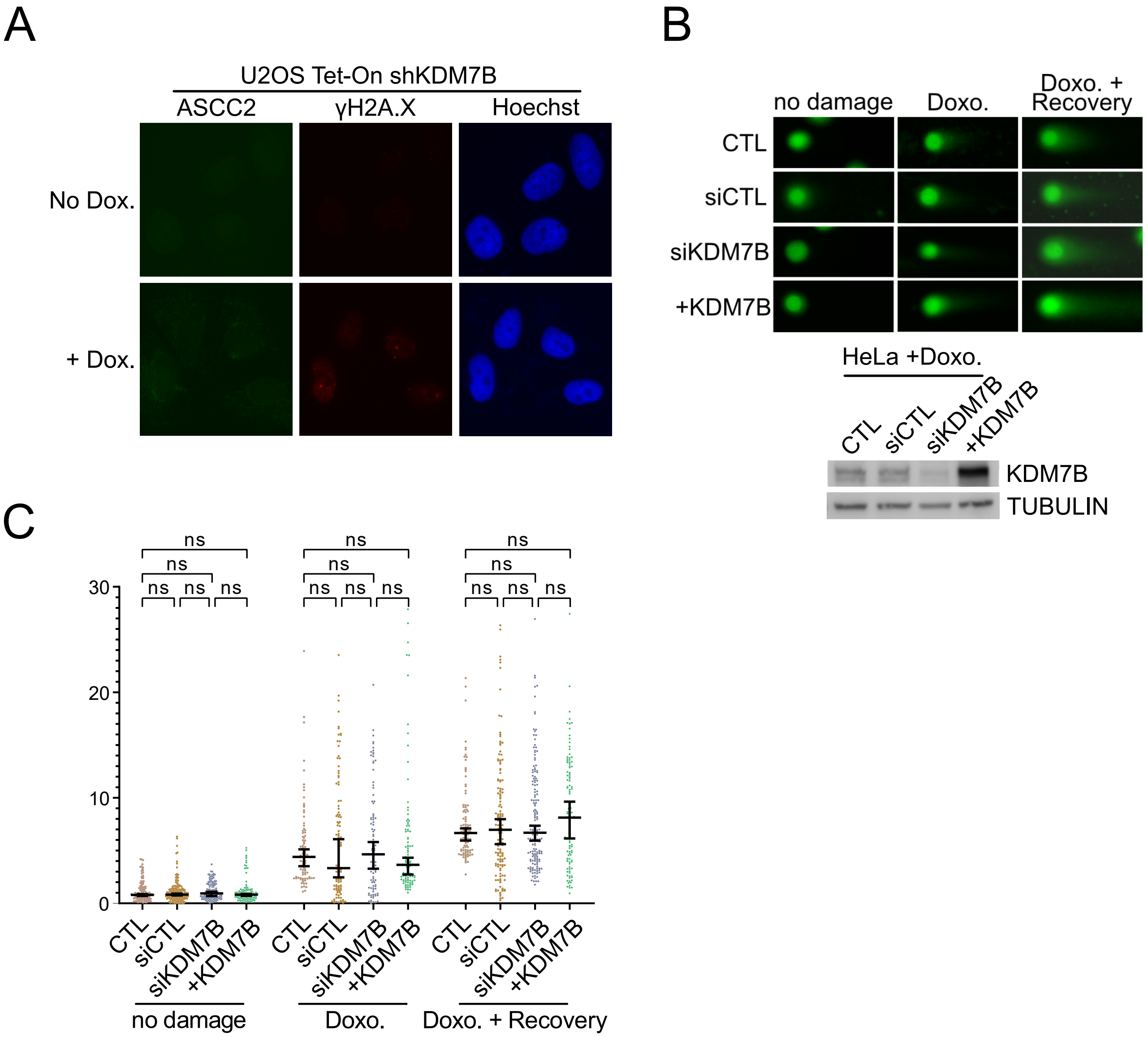

### Supplementary Figure 5

#### SUPPLEMENTARY FIGURE 5

A

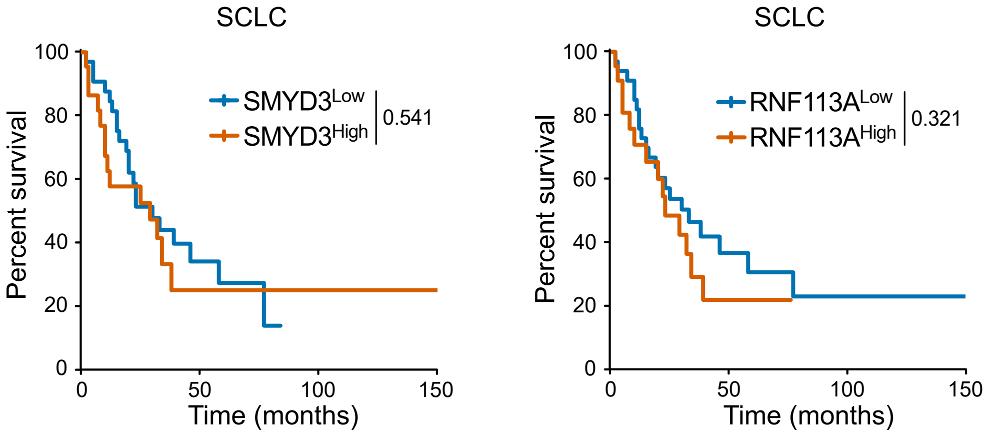


B

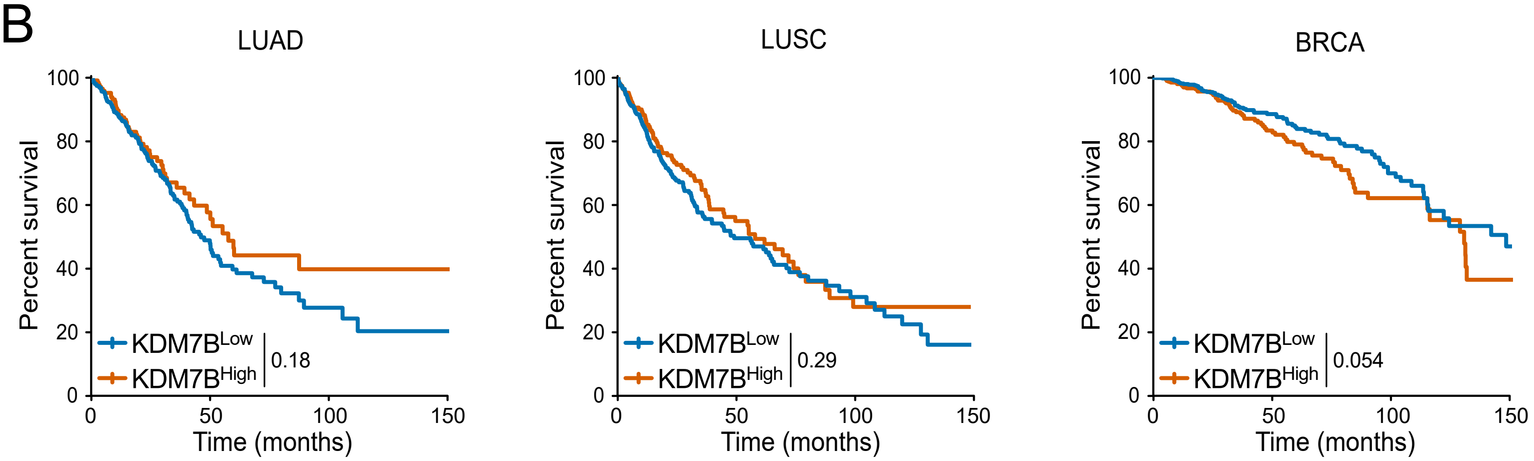

### Supplementary Figure 6

#### SUPPLEMENTARY FIGURE 6

A

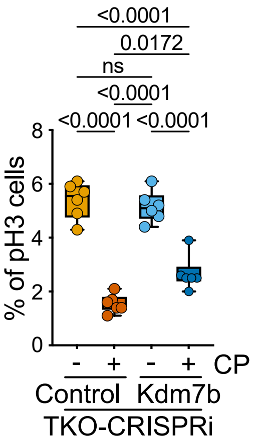


B

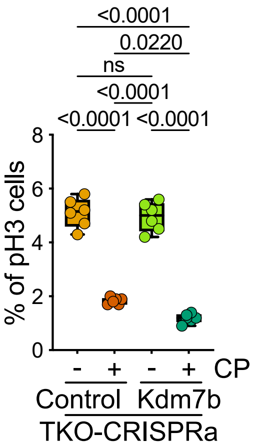
